# Mapping of the autophagosomal degradome identifies IL-7Rα as key cargo in proliferating CD4+ T-cells

**DOI:** 10.1101/2021.12.08.471825

**Authors:** Dingxi Zhou, Mariana Borsa, Daniel J. Puleston, Susanne Zellner, Jesusa Capera, Sharon Sanderson, Luke Jostins, Christian Behrends, Ghada Alsaleh, Anna Katharina Simon

**Affiliations:** The Kennedy Institute of Rheumatology, NDORMS, University of Oxford, Roosevelt Drive, OX3 7FY, Oxford, UK; Munich Cluster for Systems Neurology (SyNergy), Medical Faculty, Ludwig-Maximilians-University München, Feodor-Lynen Strasse 17, 81377, Munich, Germany

## Abstract

CD4+ T cells orchestrate both humoral and cytotoxic immune responses. While it is known that CD4+ T cell proliferation relies on autophagy, direct identification of the autophagosomal cargo involved is still missing. Here, we created a transgenic mouse model, which, for the first time, enables us to directly map the proteinaceous content of autophagosomes in any primary cell by LC3 proximity labelling. IL-7Rα, a cytokine receptor mostly found in naïve and memory T cells, was reproducibly detected in autophagosomes of activated CD4+ T cells. Consistently, CD4+ T cells lacking autophagy showed increased IL-7Rα surface expression, while no defect in internalisation was observed. Mechanistically, excessive surface IL-7Rα sequestrates the common gamma chain, impairing the IL-2R assembly and downstream signalling crucial for T cell proliferation. This study provides proof-of-principle that key autophagy substrates can be reliably identified with this model to help mechanistically unravel autophagy’s contribution to healthy physiology and disease.

## Introduction

Macroautophagy, hereafter as autophagy, is an evolutionarily conserved catabolic pathway that mediates the degradation of cellular components. Autophagy plays an essential role in the differentiation, homeostasis and renewal of many immune cells^11^. Among these immune populations is the CD4+ T cell, or helper T cell, which orchestrates both adaptive and innate immunity. To exert their helper function, CD4+ T cells need to be activated and differentiated into effector populations. The activity is finely tuned by three sequential signals: 1) T cell receptor (TCR) recognising their cognate antigen in a major histocompatibility complex (MHC)-restricted manner, 2) costimulatory receptors activated by their ligands, and 3) cytokine-amplified differentiation and expansion^12^. Moreover, activated T cells undergo a dramatic shift in their proteome, which facilitates their functional and metabolic transition^13^. Autophagy has been shown to contribute to the expansion, differentiation, and maintenance of CD4+ T cells, suggesting that autophagy can selectively degrade a spectrum of molecules to mediate these physiological processes^14–16^. The systematic identification of autophagosomal cargoes in CD4+ T cells will fill a gap in our understanding of the molecular mechanisms of T cell activation and proliferation.

Traditional methods for mapping autophagosomal content are based on purifying autophagosomes with extensive cell fractionation and mass-spectrometry analysis. However, the results are compromised by contamination from other co-purified cellular compartments and by the low reproducibility caused by variation introduced during the differential centrifugation and extensive manipulation. Furthermore, the number of cells required often exceeds what can be obtained from primary material. To determine bona fide autophagosomal constituents, the proximity-dependent biotinylation approach provides a solution. It uses a modified ascorbate peroxidase (APEX2) to transform biotin-phenols into short-lived reactive radicals, which can covalently tag proteins in close proximity^17^. When genetically fused to a protein of interest, the enzyme provides a snapshot of the local environment and maps the spatial proteome of subcellular compartments^18^. Le Guerroué et al. employed the technique to profile potential autophagosomal substrate proteins in HeLa cells through over expressing APEX2 fused with hATG8^8^. To extend its application to primary cells under different physiological contexts, we created a knock-in mouse model by introducing the APEX2 sequence into the endogenous *Lc3b* locus, an *Atg8* homolog in mammalian cells. This model helped us to identify IL-7Rα as a major autophagy cargo in proliferating CD4+ T cells, thus revealing how autophagy contributes to CD4+ T cell expansion by modulating signal 3.

## Results

### Autophagy-deficient CD4+ T cells show delayed proliferation

In a study by Ahmed et al. and our previous study, it was revealed that autophagy is crucial for the formation of memory CD8+ T cells4,19. We used a mouse model conditionally deleting *Atg7*, a key autophagy gene, in both CD4 and CD8 T lymphocytes (CD4^cre^ *Atg7^flox/flox^,* hereafter as T-Atg7^−/−^). Using the same mouse model here, we addressed the role of autophagy in CD4+ T cells and found that the formation of CD4+ effector T cells was severely affected in two models of antigen exposure (Fig. 1a-d). We excluded the effects of autophagy deletion in antigen-presenting cells expressing CD4 by generating chimera with bone marrow from wild-type littermate controls (CD45.1+) and T-Atg7^−/−^ mice (CD45.2+) (Fig. 1a). Hosts were challenged with murine cytomegalovirus (MCMV) after 7 weeks and MCMV-specific CD4+ T cells in both spleens and lungs were evaluated with MHC class II tetramers, on day 7-9, the peak time for effector T cell formation^20^. The number of CD45.2+ MCMV-specific T-Atg7^−/−^ cells were significantly lower than that of CD45.1+ wild-type cells (spleens in Fig. 1b and lungs in supplementary Fig. 1a).

**Figure 1.**
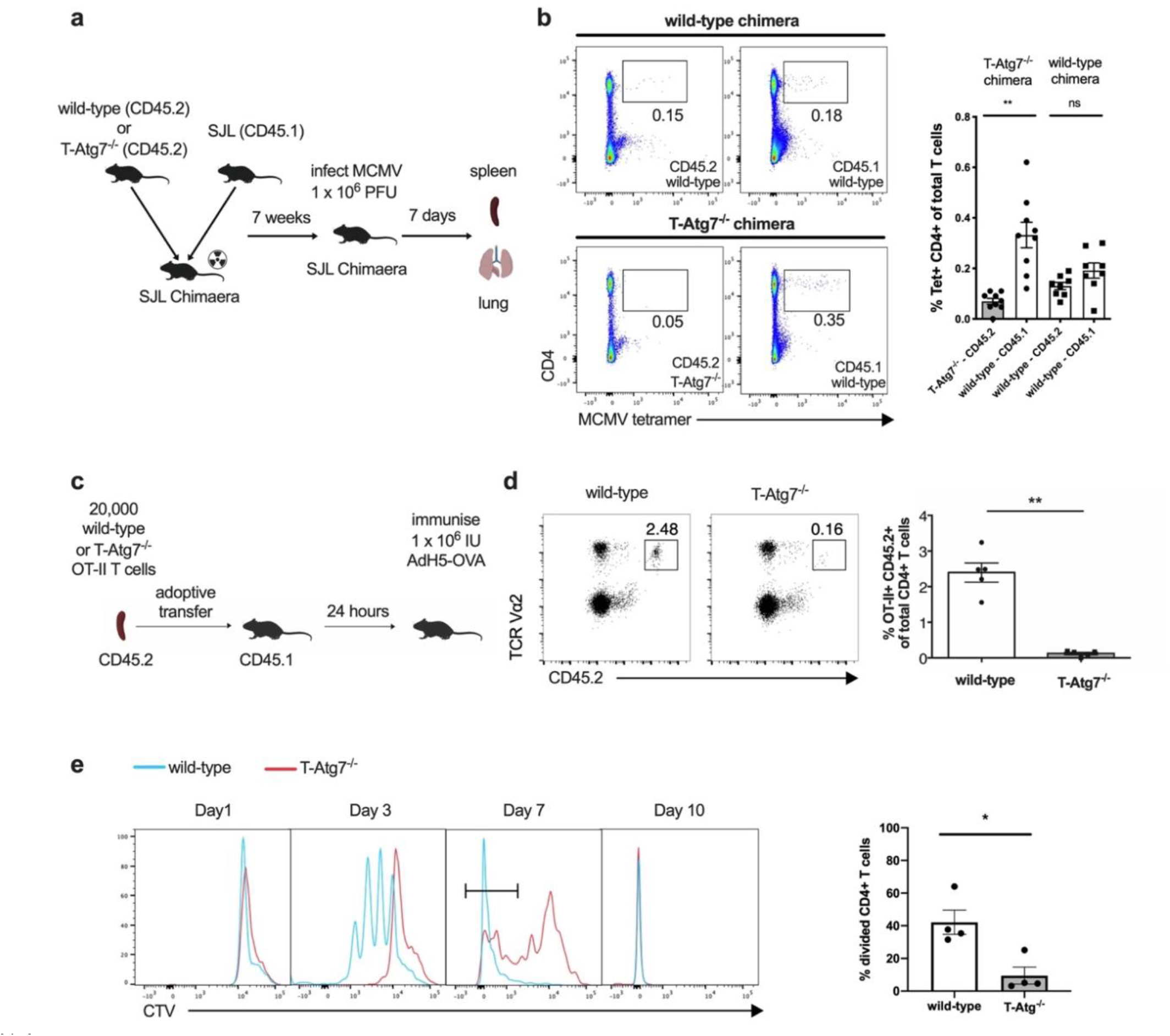
Autophagy-deficient CD4+ T cells show delayed proliferation. **(a)** Experimental set-up for the generation of bone marrow (BM) chimeras. Lethally irradiated CD45.1+ hosts reconstituted with a 1:1 mix of BM of either T-Atg7^−/−^ or wild-type (donors both CD45.2+) and CD45.1+ wild-type BM. After 7 weeks, hosts were intravenously (i.v.) infected with 1×10^6^ PFU murine cytomegalovirus (MCMV). Cells from spleens were collected and analyse 7 days post-infection. **(b)** Dot plots (left) of splenic MCMV-specific CD4+ T cells within CD19-population from CD45.2+ wild-type or T-Atg7^−/−^ donors and CD45.1+ donors. Bar graph (right) indicates percentage of Tet+ CD4+ T cells within the CD45.2+ CD19- or CD45.1+ CD19-population (n = 8-9 mice per group) and is representative of three independent experiments. **(c)** Experimental set-up for the adoptive transfer: OT-II lymphocytes (2 × 10^4^) from either CD45.2 wild-type or T-Atg7^−/−^ mice were adoptively transferred into CD45.1 recipient mice. The recipient mice were orally immunised with 1×10^6^ IU recombinant H5 adenovirus expressing ovalbumin (AdH5-OVA). The number of OT-II cells was determined by flow cytometry 8 days after immunisation. **(d)** Dot plots (left) are gated on CD45.2+, TCR Vα2+ in the spleens of recipient mice. Bar graph (right) depicts the frequency of gated population within total CD4^+^ T cell population (n = 5 mice per group) and is representative of three independent experiments. **(e)** Splenic OT-II+ T cells from wild-type or T-Atg7^−/−^ mice were stimulated with ovalbumin in culture medium supplemented with mIL2 for 1, 3, 7 or 10 days. Histograms by flow cytometry (left) represent OT-II+ CD4^+^ T cell proliferation. Bar graph (right) indicates the percentage of CellTrace Violet-negative cells (most divided, defined by gate on day 7) within total OT-II+ CD4^+^ T cells (n = 4 mice per group). Data are representative of 3 independent experiments. All data except for (b) are represented as mean ± SEM with unpaired two-tailed Student’s t-test, p*<0.05, p**<0.01, ns: not significant.

To further confirm that autophagy-deficient CD4+ T cells fail to expand upon antigenic stimulation, we bred the T-Atg7^−/−^ mice with the OT-II T-cell receptor transgenic mouse model, where T cells specifically recognise an epitope spanning peptide residues 329– 337 of ovalbumin (OVA)^21^. Neither CD4+ T cell lymphopenia nor increase of effector memory population (CD62L^−^ CD44^high^) were observed in the spleens of naïve 6-week-old T-Atg7^−/−^ OT-II mice (supplementary Fig. 1b-c). We transferred either wild-type or T-Atg7^−/−^ OT-II T cells of CD45.2+ background into CD45.1+ host mice. After 24 hours, mice were challenged with recombinant H5 adenovirus expressing OVA (AdH5-OVA) to activate the OT-II T cells (Fig. 1c). On day 9, phenotyping of antigen-specific cells revealed that autophagy-deficiency significantly compromised effector T cell formation in both spleen and blood (Fig. 1d and supplementary Fig. 1d), which is consistent with our findings measuring endogenous T cell responses to MCMV.

To understand why antigen-specific CD4+ T cells show impaired expansion in vivo, we next dissected the dynamics of the TCR-mediated proliferation in autophagy-deficient cells. We activated wild-type and autophagy-deficient OT-II CD4+ T cells in vitro and assessed their proliferation profile. We observed a delay in proliferation of T-Atg7^−/−^ CD4+ T cells compared to their wild-type counterparts on day 3 and day 7 (Fig. 1e). However, by day 10, autophagy-deficient CD4+ T cells caught up and exhibited a similar profile to wild-type. In line with previous work, the upregulation of early activation markers is not significantly changed between autophagy-deficient and - proficient cells, indicating that the TCR-signalling is not affected (Supplementary Fig. 1e and f)^22^.

To rule out the preferential proliferation (and escape) of CD4+ T cells with incomplete *Atg7* knockout, we tested *Atg7* mRNA expression by qRT-PCR on day 7 in sorted T cells that had undergone a maximum number of detectable cell divisions (i.e. that diluted the cell dye to the maximum). Indeed, *Atg7* mRNA in proliferated CD4+ T cells from T-Atg7^−/−^; OT-II mice was undetectable (Supplementary Fig. 1g). Moreover, decreased expansion cannot be explained by increased apoptosis or cell death, since the percentages of Annexin V+/LD-NIR- and LD-NIR+ CD4+ T cells do not differ significantly between wild-type and knockout cells on day 1 or 3 (supplementary Fig. 1h and i). Our observation of delayed proliferation in response to antigen-stimulation confirms and extends the findings of other autophagy-knockout models that demonstrated an issue with CD4 expansion in response to non-specific stimulation^22–26^. Next, we aimed to identify the proteins that are selectively degraded by autophagy by using proximity labelling.

### A novel mouse model based on the proximity biotinylation technique

Among the six mammalian ATG8 homologs, MAP1LC3B (LC3B) is the most widely used to detect autophagic flux. Furthermore, in the study by Le Guerroué et al., autophagosomal substrates labelled by LC3B-fused APEX2 (AP2) displayed the highest reproducibility between biological samples among all six ATG8 homologs8. Therefore, we created a novel transgenic mouse model, knocking the sequence of AP2 and a flexible GS-linker into the endogenous *Lc3b* locus (Fig. 2a). In the presence of both biotin-phenol (BP) and hydroxyl-peroxide (H2O2), LC3B-fused AP2 transforms biotin-phenol into highly reactive radicals, which can further be covalently conjugated to proteins nearby (Figure 2a). Then, these biotinylated proteins undergo affinity purification and mass spectrometry-based profiling. LC3B is expressed on the cytosolic and luminal side of autophagosomes but only on the cytosolic side of LC3-associated phagosomes27,28. To increase the chance of discovering targets on the luminal side of autophagosomes, homogenates extracted from the proximity-labelled cells, which contains intact autophagosomes and other organelles, were incubated with Proteinase K (Prot K) (Fig. 2a). This allows Prot K to degrade the cytosolic proteins bound to LC3B, while those residing inside autophagosomes and other vesicles remained largely intact due to the protective effect of the membranes9.

**Figure 2.**
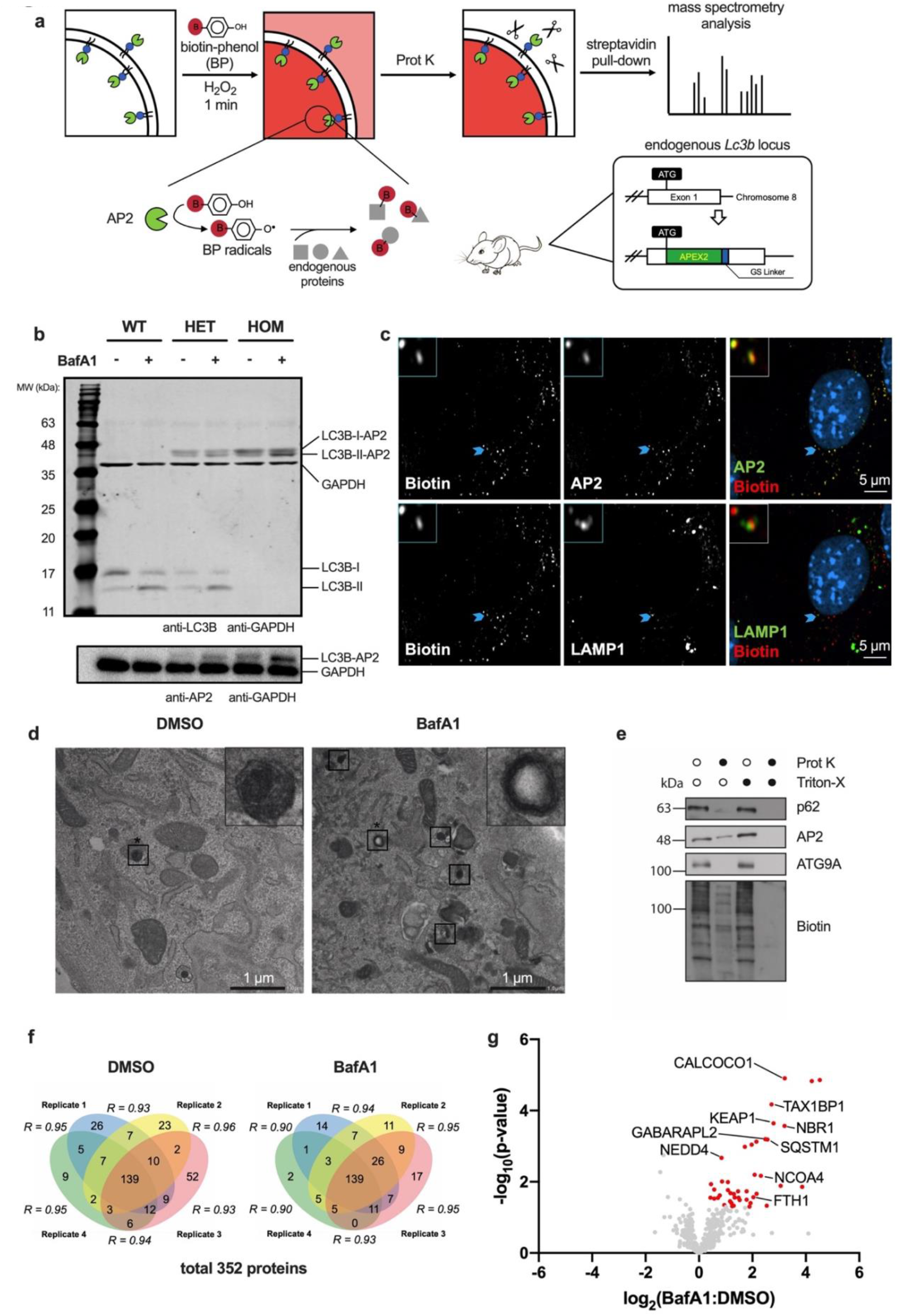
LC3B-AP2 transgenic mouse model facilitates the direct identification of autophagosomal cargoes. **(a)** Generation of mouse model and experimental set-up for proximity labelling. **(b)** Western blot of whole splenic cell homogenates from wild-type, heterozygous and homozygous mice. Cells were treated with 10 nM Bafilomycin A1 (BafA1) or DMSO for 2 hours. These experiments were repeated as biological triplicates (one mouse each) with similar results. **(c)** Confocal imaging of biotinylated proteins, AP2 and LAMP1 in proximity-labelled immortalised mouse embryonic fibroblasts (imMEFs) generated from the *Lc3b-AP2* mouse model. Light blue arrows indicate colocalization events with magnifications at the upper-left corners. These experiments were repeated as biological triplicates with similar results. **(d)** Electron micrographs of DMSO- or BafA1-treated *Lc3b-AP2* imMEFs. Following fixation, cells were incubated with 3,3-diaminobenzidine and H_2_O_2_ prior to standard embedding and ultrathin sectioning. Squares highlight the autophagosomal structures. Squares that are marked by asterisks are magnified in the upper-right corner. **(e)** Homogenates from imMEFs were labelled, then left untreated or incubated with Prot K, Triton-X-100, or both and followed by immunoblotting for AP2, Biotin, p62 and ATG9A. **(f)** Venn diagrams depicting Prot K-protected, biotinylated proteins identified in imMEFs with more than two spectral counts across four biological replicates in both DMSO and BafA1 treated groups. Pearson’s correlation coefficients of the label-free quantity of each protein included in two replicates are displayed in italic. **(g)** Volcano plot of proteins from (f) labelled by LC3B-AP2 in imMEFs. Proteins significantly upregulated in response to BafA1 treatment are highlighted in red (p<= 0.05 by unpaired two-tail Student’s t-test, n = 4 biological replicates). Known autophagosomal candidates are labelled with protein names.

Wild-type, heterozygous and homozygous mice are identified by genomic PCR (supplementary Fig 2a). We first confirmed that the LC3B-AP2 chimeric protein is expressed. Cellular LC3B has two major isoforms – the free LC3B-I located in the cytosol and the membrane-bound LC3B-II. Immunoblotting of splenocytes showed bands around 44 kDa in heterozygotes and homozygotes with both anti-LC3B (17 kDa) and anti-AP2 (27 kDa) staining, but not in wild-type cells (Fig. 2b). In wild-type and heterozygous splenocytes, LC3B was mostly expressed as non-fused to AP2. As expected, the lipid-conjugated form of LC3B-AP2 (LC3B-II-AP2) and non-fused LC3B-II accumulated when cells were treated with Bafilomycin A1 (BafA1), an inhibitor preventing the lysosomal degradation of LC3 in autophagosomes. These results confirm that LC3B-fused AP2 is targeted to autophagosomes and undergoes lysosomal degradation, as expected from a functional LC3B protein. Next, we determined the enzymatic activity of AP2. After 30 min BP pre-incubation and 1 min H_2_O_2_ treatment, mouse splenocytes were lysed and analysed by western blotting with streptavidin. Cellular lysates from homozygote mice (two copies of *Lc3b-AP2*) treated with BP and H_2_O_2_ showed the strongest bands spanning from 11 kDa to 200 kDa, whereas negative controls in which we omitted BP or H_2_O_2_, or cell lysates from mice that do not express LC3B-AP2, showed only weak biotin bands (Supplementary Fig. 2b).

To optimise the protocol of autophagosomal protein purification, we immortalised mouse embryonic fibroblast (imMEF) extracted from *Lc3b-AP2* embryos by transfecting them with lentivirus expressing SV40 antigens. In the imMEFs, as expected, we observed the co-localisation of biotinylated proteins with AP2 and LAMP1, a lysosomal marker (Fig. 2c), further confirming the autophagosomal enrichment of LC3B-AP2. In addition, the LC3B-AP2 chimeric protein generated electron microscope-dense autophagosomal structures when cells were subjected to 3,3-diaminobenzidine labelling and H_2_O_2_ pulsing, which were more prominent with BafA1-treatment (Figure 2d). To further validate that the LC3B-AP2 chimeric protein is indeed located on the autophagosomal inner membrane, thus labelling luminal proteins, we performed Prot K protection assays. Homogenates from proximity-biotinylated imMEFs were incubated with Prot K and/or Triton-X, a detergent that breaks all membrane structures. Compared to untreated or Triton-X treated control, the sample treated with only Prot K partly preserved p62, an autophagy cargo receptor protein, AP2 and biotinylated bands, while the cytosolic tail of the transmembrane protein ATG9A was completely degraded (Fig. 2e). By contrast, adding both Prot K and Triton-X removed all proteins. Together, these results indicate the correct localisation of LC3B-AP2 proteins in autophagosomes and the protective effect of the autophagosomal membranes to Prot K.

To test whether our mouse model can indeed help identify autophagy substrates, we next performed quantitative MS-based proteomics using *Lc3b-AP2* imMEFs (Supplementary Fig. 2c). Cells were treated or not with BafA1 and four biological replicates were included in each group. After excluding Prot K-resistant candidates found when cells were treated with MS-compatible detergent RAPIGest, we reproducibly identified a total of 352 proteins specifically protected from Prot K treatment (Fig. 2f and Supplementary Table 1)^29^. Among the Prot K-protected proteins, 43 proteins were significantly increased upon BafA1 treatment (Fig. 2g). Gene ontology analysis revealed that the identified proteins belong to the category of proteins that are usually enriched in autophagosomes and other cytoplasmic vesicles, where the membrane-bound LC3B is mainly located (Supplementary Fig. 2d)^30^. As expected, most of the proteins identified under basal conditions include autophagic receptors (p62/SQSTM1, NBR1, NCOA4, CALCOCO1 and TAX1BP1), autophagy regulators interacting with LC3 (NEDD4), autophagy substrates (KEAP1, FTH1) and another ATG8 homolog (GABARAPL2). Taken together, these observations support the reliability of our mouse model, which can be used as a tool to specifically identify autophagosomal cargoes.

To determine the minimum cell number to detect autophagosomal cargo effectively, we performed a cell titration experiment with cell numbers ranging from 0.125 to 8 million. Although two autophagic receptors, TAX1BP1 and p62/SQSTM1, could be consistently identified as BafA1-sensitive candidates in all conditions, the total number of proteins being significantly upregulated by BafA1 showed a sharp decrease when less than 1 million imMEFs were used (Supplementary Fig. 2e). These data indicate that a greater cell number potentially yields more candidates when using the LC3B-AP2 labelling system.

### IL-7Rα is detected in LC3-containing vesicles in activated CD4+ T cells

Aiming to better understand the underlying causes of the observed delayed proliferation in autophagy-deficient CD4+ T cells, we decided to apply the LC3B-AP2 labelling system to investigate the autophagosomal degradome in autophagy-proficient proliferative CD4+ T cells. For that, we used activated CD4+ T cells isolated from *Lc3b-AP2* mice. The proliferative state of CD4+ T cells allowed us to obtain a large number of fully activated CD25+ cells, which, based on the results obtained from imMEFs, contributes to the identification of higher numbers of autophagosomal candidates (Supplementary Fig. 3a). In addition, we could determine that the autophagic flux was highest on days 2 and 3 (Supplementary Fig. 3b), indicating these are appropriate time points to evaluate the role of autophagy in CD4+ T cell proliferation. Finally, we identified an accumulation of LC3B-II-AP2 protein in BafA1-treated cells 3 days post-activation (Supplementary Fig. 3c). Thus, we decided to activate CD4+ T cells in vitro for 3 days, using anti-CD3/CD28 Dynabeads, prior to performing proximity labelling.

Again, the mass-spectrometry analysis showed high reproducibility between each biological replicates (Fig. 3a). Among 112 Prot K-protected and streptavidin-enriched proteins, seven candidates were significantly upregulated by BafA1-treatment (Fig. 3b and Supplementary Table 2). Of these candidates, interleukin-7 receptor alpha (IL-7Rα) showed the highest fold change robustly among all biological replicates (4.67-fold increase, p = 0.033, Fig. 3c). Moreover, IL-7Rα was found to be significantly upregulated (p = 0.0089) in a repeat experiment, and a meta-analysis combining the two experiments found that IL-7Rα was the only protein that was significant after multiple testing correction (meta-analysed p = 0.00087, Benjamini-Hochberg-corrected q-value = 0.0315, Supplementary Fig. 3d and Supplementary Table 3). Therefore, we decided to further investigate this protein. IL-7Rα is highly expressed on naive T cells, and TCR/CD28-mediated signalling induces its downregulation^31^. This is in line with our data suggesting that IL-7Rα is degraded in an LC3B+ compartment during T cell activation.

**Figure 3.**
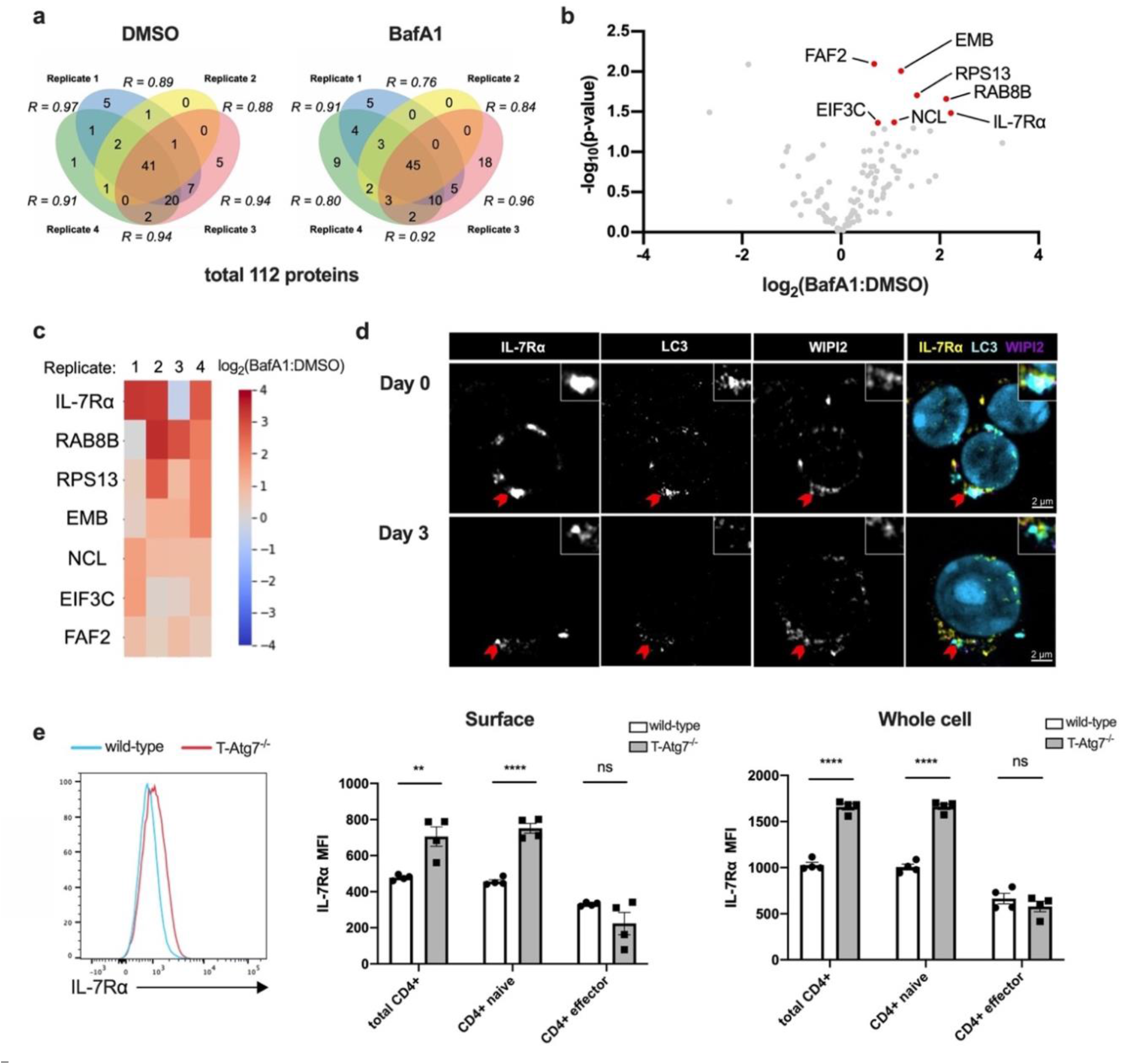
IL-7Rα is degraded via autophagy in CD4+ T cells. **(a)** Venn diagrams depicting Prot K-protected, biotinylated proteins identified in activated CD4+ T cells with more than two spectral counts across four biological replicates in both DMSO and BafA1 treated groups. Pearson’s correlation coefficients of the label-free quantity of each protein included in two replicates are displayed in italic. **(b)** Volcano plot of proteins from (a) labelled by LC3B-AP2 in activated CD4+ T cells. Proteins significantly upregulated in response to BafA1 treatment are highlighted in red, p<0.05 by unpaired two-tailed Student’s t-test, n = 4 biological replicates per group, with each replicate combining T cells from two mice. **(c)** A log2(BafA1/Ctrl) heat map of the BafA1-upregulated candidates. Red colour indicates an upregulation in BafA1-treated samples, while blue represents a decrease. **(d)** Confocal imaging of IL-7Rα, LC3 and WIPI2 in wild-type CD4+ T cells without activation or activated for 3 days. Red arrows indicate colocalization events, whose magnifications are displayed at the upper-right corners. These experiments were repeated as biological triplicates with similar results. **(e)** Histogram (left) of IL-7Rα surface level, gated on splenic CD4+ T cells from wild-type or T-Atg7^−/−^ mice. Bar graphs show the surface (middle) or whole-cell (right) level of IL-7Rα in total CD4+ T cell, naïve CD4+ T cell (CD62L^+^ CD44^lo^) or effector CD4+ T cell (CD62L^−^ CD44^hi^). Quantitative analyses are representative of four independent experiments. All values are represented as mean ± SEM with unpaired two-tailed Student’s t-test, n = 3-4 mice per group, p*<0.05, p***<0.001, p****<0.0001, ns: not significant.

IL-7Rα is a subunit of the heterodimeric cytokine receptor IL-7R, and its signalling is crucial for peripheral T cell survival and homeostatic proliferation^32^. It is constantly internalised and recycled through membrane-enveloped organelles and therefore may be found in LC3-containing compartments other than autophagosomes^33,34^. To ensure that IL-7Rα is found in bona fide autophagosomes, we performed immunostaining of IL-7Rα with LC3 or WIPI2, an early autophagosomal marker. We observed colocalization of IL-7Rα with both molecules when cells are either activated or not (Fig. 3d). Therefore, we confirmed that IL-7Rα was indeed located in the autophagosomal compartment.

### IL-7Rα accumulation in T-Atg7^−/−^ CD4+ T cells is due to impaired autophagy

To validate whether lack of autophagy impairs the degradation of IL-7Rα, we measured IL-7Rα expression level in CD4+ T cells freshly isolated from the spleen of T-Atg7^−/−^ mice by flow cytometry. IL-7Rα was upregulated both at the surface and whole-cell level in naïve T-Atg7^−/−^ CD4+ T cells (Fig. 3e). By contrast, no significant changes in effector T cells were observed. To make sure this is an autophagy-related effect and not an *Atg7*-specific effect, we further confirmed this result by deleting *Atg16l1*, another autophagy gene, specifically in T cells (Supplementary Fig. 3e). The increased expression of IL-7Rα protein was not due to enhanced transcription of *Il7ra* as indicated by the qRT-PCR performed on different days after activation (Supplementary Fig. 3f). It is worth noting that both surface and whole-cell levels of IL-7Rα remained higher in T-Atg7^−/−^ CD4+ T cells after one day of activation, while the difference disappeared on day 3 (Supplementary Fig. 3g).

Previous studies showed that key autophagy genes, like *Atg7* and *Atg5*, control the internalisation of surface receptors^5,35^. To address whether the accumulation of IL-7Rα in autophagy-knockout CD4+ T cells is caused by decreased internalisation, we measured the amount of IL-7Rα binding to the biotin-conjugated antibody at the cell surface over time. We confirmed that the surface level of IL-7Rα was generally higher in T-Atg7^−/−^ naïve CD4+ T cells in comparison to their wild-type counterparts, but could not observe any differences in the rate of receptor internalisation, represented by the percentage of internally trafficked IL-7Rα (Supplementary Fig. 3h). Therefore, we ruled out the possibility that the surface accumulation of IL-7Rα is due to impaired internalisation.

### Excessive IL-7Rα on autophagy-deficient T cell surface impair the IL-2R assembly and signalling

While we confirmed that the accumulation of IL-7Rα in CD4+ T cells is autophagy-dependent, the molecular basis of how it leads to impaired proliferation remained elusive. Both IL-7R and IL-2R share a common gamma-chain (γc) to transduce downstream signalling^36^. IL-7R (heterodimer comprised of γc and IL-7Rα) is highly expressed on naïve T cells and co-exists with the low-affinity IL-2R (heterodimer comprised of γc and IL-2Rβ). During TCR-mediated activation, IL-7Rα is downregulated from the cell surface and IL-2Rα takes over to form high-affinity IL-2R (heterotrimer comprised of γc, IL-2Rα and IL-2Rβ). When binding to IL-2, high-affinity IL-2R mediates persistent and robust downstream signalling, which is crucial for the proliferation of CD4+ T cells^37^. It has been reported in regulatory T cells that γc has limited availability and pre-associates with excessive IL-7Rα independently of IL-7, which impedes IL-2R signalling^38^. Therefore, we hypothesized that excessive IL-7Rα expression leads to delayed proliferation of T-Atg7^−/−^ CD4+ T cells by a similar mechanism (Fig. 4a).

**Figure 4.**
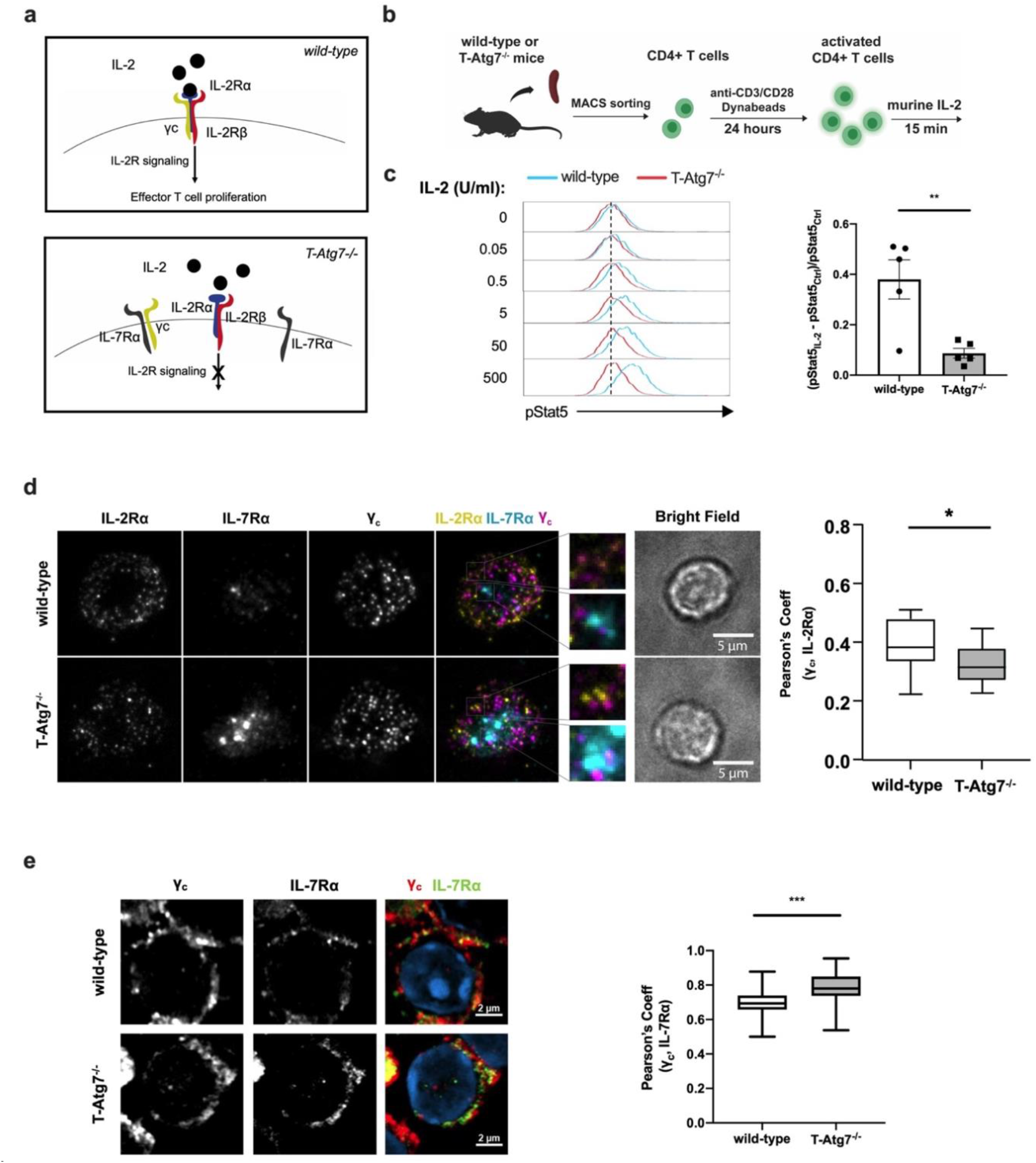
IL-2R signalling is impaired in autophagy-deficient CD4+ T cells. **(a)** Schematic overview of how excessive IL-7Rα may lead to impaired T cell proliferation. During T cell activation, the IL-2Rα is upregulated on the cell surface, while surface IL-7Rα is decreased. In T-Atg7^−/−^ CD4+ T cells, excessive IL-7Rα sequestrates the common gamma chain (γc), a key subunit of both IL-7R and IL-2R, and abrogates the downstream IL-2R signalling which is crucial for the expansion of the effector population. **(b)** Experimental set-up: naïve CD4+ T cells from either wild-type or T-Atg7^−/−^ mice were activated with anti-CD3/CD28 Dynabeads for 24 hours, followed by a treatment with murine IL-2 for 15 min. Phosphorylated Stat5 (pStat5) level was measured by flow cytometry. **(c)** Histograms (left) of pStat5 level in activated CD4+ T cells from wild-type or T-Atg7^−/−^ mice, stimulated with different doses of murine IL-2. Bar graph (right) depicts the ratio of increase in p-Stat5 signal in response to 20 ng/ml murine IL-2 (pStat5_IL-2_ – pStat5_ctrl_) to the basal level (pStat5_ctrl_), determined from two independent experiments, n = 5 mice per group. Data are represented as mean ± SEM with unpaired two-tailed Student’s t-test, p**<0.01. **(d)** TIRF imaging (left) of IL-7Rα, IL-2Rα and γc at the immunological synapse. Naive CD4+ T cells activated for 24 hours were exposed to supported lipid bilayers containing ICAM1 and anti-CD3 proteins, then fixed after a 10-min incubation with 50 U/ml mIL-2. Cytokine receptors were stained for imaging. Square insets show magnification of the co-localisation events. Box graph (right) shows Pearson’s correlation coefficient between the IL-2Rα and γc, n = 15 cells per group. Data are represented as mean ± max and min values, with unpaired two-tailed Student’s t-test, p*<0.05. **(e)** Confocal imaging of IL-7Rα and γc at the surface of activated CD4+ T cells (with high CD25 expression). Box graph (right) shows Pearson’s correlation coefficient between the IL-7Rα and γc at the cell surface, n = 23 cells per group. Data are represented as mean ± max and min values, with unpaired two-tailed Student’s t-test, p***<0.001.

To test whether IL-2R signalling is impaired in CD4+ T cells lacking autophagy, we measured the level of phosphorylated Stat5 (pStat5), the classical downstream transducer of IL-2 signalling. Since naïve T cells respond poorly to IL-2, we hypothesized that IL-2 signalling contributes to T cell activation mainly after high-affinity IL-2R is expressed^39^. We observed an upregulated expression of IL-2Rα 24 hours after TCR-mediated activation (Supplementary Fig. 3a), while IL-7Rα remained accumulated on the cell surface of autophagy-deficient CD4+ T cells at the same time point (Supplementary Fig. 4a). To determine whether accumulated IL-7Rα impacts on IL-2-driven signalling, we incubated activated CD4+ T cells with different doses of IL-2 (Fig. 4b). Corroborating our initial hypothesis, pStat5 level was higher in presence of increased concentrations of IL-2 in wild-type CD4+ T cells, whereas IL-2R signalling in T-Atg7^−/−^ T cells was refractory to the administration of IL-2 (Fig. 4c). Importantly, this was not due to different expression of IL-2R, as we found no significant differences in the expression of its three subunits between T-Atg7^−/−^ and wild-type T cells (Supplementary Fig. 4b).

To further confirm whether the assembly of these cytokine receptors is impaired in T-Atg7^−/−^ CD4+ T cells, we measured the co-localisation of γc with either IL-7Rα or IL-2Rα. Since IL-2 signalling is polarised at the immunological synapse, we incubated the activated T cells on supported lipid bilayers containing ICAM1 and anti-CD3 proteins, in the presence of IL-2^40^. We then visualised the receptor subunits at the immunological synapse (Fig. 4d, left). As expected, we found a significantly lower co-localisation of IL-2Rα and γc in T-Atg7^−/−^ CD4+ T cells (Fig. 4d, right), indicating that the formation of IL-2R complex was indeed impaired in autophagy-deficient CD4+ T cells at the immunological synapse. However, we did not see any significant differences in the colocalization of γc with IL-7Rα (Supplementary Fig. 4b). To evaluate whether IL-7Rα sequestration happened outside the immunological synapse, we performed confocal analysis of whole cell volumes, where we could observe that γc-IL-7Rα co-localisation is indeed higher in T-Atg7^−/−^ CD4+ T cells (Fig. 4e). Taken together, our results suggest that autophagy plays a key role in CD4+ T cell activation, by degrading IL-7Rα, which releases γc and promotes its assembly in high-affinity IL-2R, a signalling receptor that is essential for T cell proliferation.

## Discussion

In this study, we dissected the molecular mechanism of autophagy in the antigen-induced proliferation of CD4+ T cells. Using a novel powerful mouse model based on the proximity biotinylation technique, we demonstrate that autophagic degradation of IL-7Rα plays a crucial role in T cell activation, since it can competitively inhibit IL-2R signalling through sequestration of the limited amount of γc. Thus, we identified autophagy’s role in the transition of cytokine signalling, from IL-7R-mediated homeostasis in naïve T cells to IL-2R-mediated expansion in effector T cells. This research not only provides proof-of-principle application of the *Lc3b-AP2* mouse model but also fills a key gap in our knowledge of CD4+ T cell activation.

We observed that IL-7Rα levels were higher in autophagy-deficient naïve CD4+ T cells, while the difference disappears on 3 days post-activation, suggesting that other compensatory pathways for degradation of excessive IL-7Rα might take place. Previous studies indicated that IL-7Rα can be degraded through both lysosome- and proteasome-mediated pathways in naïve T cells^33,34^. During T cell activation, genes of proteasome subunits are upregulated, which may contribute to the compensatory degradation of IL-7Rα in autophagy-deficient CD4+ T cells^41^. Other pathways compensating gradually for the loss of autophagy in T-Atg7^−/−^ CD4+ T cells may help to explain why the cells catch up on their proliferation in vitro.

Our observation of accumulated IL-7Rα in both *Atg7*- and *Atg16l1*-deficient CD4+ T cells is in contrast to a report in T cells lacking Vps34, a PI3-kinase involved in the initiation of autophagy^42^. In this study, McLeod et al. describe that Vps34-deficient CD4+ T cells downregulate the surface expression of IL-7Rα. However, the autophagic flux in that model is not impaired and the lower level of surface IL-7Rα was proposed to be due to a deficit in the retromer pathway mediated by Vps34. By contrast, in another study in which bone marrow chimera model were generated to exclude the effect of lymphopenia, IL-7Rα expression is higher in Vps34-deficient CD4+ T cells compared to autophagy-proficient cells, further corroborates our findings, which were also made in a non-lymphopenic environment^43^.

It is known that proteins firstly described for their function in autophagy can perform diverse other functions^44^. Indeed, the autophagy machinery also participates in cellular processes other than classical macroautophagy, such as endosomal microautophagy (eMA), and in particular, LC3-associated phagocytosis (LAP) and LC3-associated endocytosis (LANDO), where lipidated LC3 is expressed on the cytosolic surface of vesicles^30^. However, any proteins associated with these vesicles would be excluded from our analysis, as LC3 would be located towards the cytosol, and our protocol specifically enriches for autophagosomal luminal proteins. Another pathway to be considered is the LC3-dependent extracellular vesicle loading and secretion (LDELS), where lipid-conjugated LC3s reside in the lumen of extracellular vesicles^45^. In fact, using a pulse-chase proximity labelling strategy, the LC3-dependent secretome of extracellular vesicles was recently profiled in vitro^45^. Interestingly, we found a putative target potentially involved in this pathway. Rab8b, a small Rab GTPase indispensable for exocytic trafficking of post-Golgi vesicles to the plasma membrane was revealed as a putative cargo in the LC3 compartment in activated CD4+ T cells (Fig. 3b and c)^46^. In addition to this role, the protein has also been suggested to be co-localised with LC3 to mediate autophagy-based secretion and autophagosome maturation^47,48^. In summary, since membrane-bound LC3 is prevalent in degradative and non-degradative vesicles, the *Lc3b-AP2* mouse model is not only a powerful technique to reveal macroautophagic cargo but may also identify other cargoes when BafA1 is replaced with other specific inhibitors. In addition, this model will be useful to explore the cytosolic subproteome of LC3 under physiological conditions when the Prot K treatment is left out.

Two other molecules, CDKN1B/p27Kip, a cell cyclin-dependent kinase inhibitor, and PTPN1, a protein tyrosine phosphatase, were shown to be removed by autophagy during the early activation of naïve CD4+ T cells and Th1 cells respectively^49,50^. CDKN1B/p27Kip1 was found upregulated by Jia et al. using whole-cell proteomics of autophagy-knockout T cells^49^. Further validation showed that the autophagy-mediated degradation of CDKN1B is a prerequisite for the proliferation of naïve CD4+ T cells. Proteomics and western blots have been widely used to indirectly map autophagosomal content from autophagy-deficient T cells. However, these techniques cannot exclude that cells without autophagy may have synthesized these as new proteins. By purifying autophagosomes with extensive subfractionation from activated Th1 cells, Mocholi et al. showed that autophagy degrades PTPN1^50^. It is possible that we did not identify either of these proteins due to the duration of activation. Both studies used T cells activated for no longer than 24 hours, while we activated cells for three days to maximise autophagy levels and cell numbers.

The *Lc3b-AP2* mouse model overcomes the obstacles of previous methods and facilitates the identification of transient binding partners and subcellular structures. One caveat of the method is the biased detection of the proteome. Proteins embedded in large complexes, or with very few electron-rich amino acid residues exposed at the surface, have a lower chance to be tagged by the biotin-phenol radicals, which limits the use of this method to detect a certain type of proteins. In our study activated primary CD4 T cells show greater biological and technical variation than imMEFs. This is most likely due to the high amount of systemic variation in immune development and variation induced by activation/ expansion in T cells, which made this pilot study particularly challenging. In addition, T cells are smaller in size with little cytoplasm compared to imMEFs. As a result, a large number of T cells were required to map autophagy-regulated proteins. However, this pilot study has proven that it is possible to detect abundant proteins (such as IL-7Rα) targeted for autophagic degradation in T cells. Identifying less abundant proteins targeted by the autophagosome in T cells will require further optimisation and more repeats. This is not necessarily true for other primary cell types. Therefore, our model, together with other murine models based on this technique, indicates their broader application for the study of proteins in organelles in primary cells and in vivo^51–56^.

Overall, it is likely that autophagy is required to selectively degrade a spectrum of molecules at different time points to lift the brake on T cell proliferation, which is finely tuned by multiple signalling pathways. An autophagic programme will also be key to remodelling the cell and providing energy, nutrients and building blocks for protein synthesis and for the generation of the daughter cells.

## Materials and Methods

### Mice

*Atg7^flox/flox^* mice1 (from M Komatsu) and CD4^cre^ mice^2^ (from Adeline Hajjar) were crossed to obtain CD4^cre^ *Atg7^−/−^* mice (T-Atg7^−/−^) on a C57BL/6 background. All mice were 6-8 weeks of age at the start of each experiment and were matched in age and gender. CD4^cre^ *Atg7+/+* littermates were used as wild-type controls (wild-type). C57BL/6 SJL CD45.1 mice for bone marrow chimera and T cell adoptive transfer were purchased from Charles River, UK. OT-II mice were crossed with CD4^cre^ *Atg7-/*-mice. CD4^cre^ *Atg16l1−/−* mice were from Kevin Maloy. *Lc3b-APEX2* (*Lc3b-AP2*) mice were generated by Cyagen, by constitutively knocking in an APEX-tag and a GGGGSGGGGGS-linker into the exon 1 of the mouse *Map1lc3b* allele. All mice were held in BSU at the Kennedy Institute of Rheumatology. All mice were under specific pathogen-free level maintenance at 24 °C, 50% humidity, and a 12:12 h light/dark cycle. Mice were kept in individually ventilated cages, with ad libitum access to autoclaved water and irradiated food pellets.

### Viral vectors and lentivirus production

P-Lenti CMV/TO SV40 small + Large T (w612-1) was a gift from Eric Campeau (Addgene plasmid # 22298). For packaging the virus, HEK293T cells were transfected with vector and replication-incompetent lentiviral packaging constructs, harvested after 48-hour post-transfection, and viral supernatants filtered through a 0.45 μM cellulose acetate filter.

### Cell line generation

The mouse embryonic fibroblasts (MEFs) were generated according to published protocols^3^. In brief, *Lc3b-AP2* homozygous embryos were separated from female uteri on E13.5. After removing the head above eyes and red organs (heart and liver), the rest of the embryos were minced and subjected to 0.25% Trypsin (Sigma) digestion, with intermittent pipetting up and down. Digestion was terminated by MEF culture media (Dulbecco’s Modified Eagle Medium, Sigma-Aldrich) supplemented with 10% Fetal Bovine Serum (Sigma-Aldrich), 2% penicillin-streptomycin (Sigma-Aldrich) and 2 mM L-Glutamine (Sigma-Aldrich). Single cells and clusters were transferred onto the T25 flask and incubated with MEF culture media overnight. After replacing the old media and removing the debris, cells were passaged until reaching confluency. To immortalise the *Lc3b-AP2* cell line, MEFs were re-seeded to 6-well plates and transduced with the supernatant of the lentivirus expressing SV40 small and large T antigens overnight. Monoclonal immortalised MEFs were selected through dilution cloning and over 10 rounds of passaging.

### Bone marrow (BM) chimera

Bone marrow chimeras were generated similarly as described before^4^. After erythrocyte lysis, BM cells extracted from a single 8-week-old wild-type or T-Atg7^−/−^ (both CD45.2) were 1:1 mixed with C57BL/6 SJL mouse (CD45.1) in a total volume of 200 μl PBS, with 3 × 10^6^ cells each. The mixture was injected intravenously into C57BL/6 SJL CD45.1 recipients 2 hours after being lethally irradiated (450 cGy twice, 4 hours apart). Followed by an 8-week reconstitution, BM chimeras were immunised with 1 × 10^6^ PFU MCMV (Smith strain ATCC:VR194).

### T-cell adoptive transfer

For adoptive transfer, 2 × 10^4^ purified splenic OT-II T cells from CD45.2 wild-type or T-Atg7^−/−^ mice were intravenously injected into C57BL/6 SJL CD45.1 recipients. After 24 hours, recipient mice were immunised with 1×10^6^ IU recombinant adenovirus H5-ovalbumin (abm).

### CD4+ T Cell isolation

Total CD4+ T cells were purified with EasySep Mouse T Cell Isolation Kit (Stemcell Technology). Total splenocytes suspended in MACS buffer (2% FBS in DPBS) were incubated with rat serum and isolation cocktail for 10 min at room temperature, then incubated with Streptavidin RapidSpheres for 2.5 min. After topped up to 2.5 ml, tubes were placed into the magnet and incubated at room temperature for 2.5 min. Purified CD4+ T cells were transferred to new tubes for further analysis.

### CD4+ T cell in vitro activation

T cells were cultured in RPMI-1640 Medium (Sigma-Aldrich) containing 10% Fetal Bovine Serum (Sigma-Aldrich), 2% penicillin-streptomycin (Sigma-Aldrich) and 2 mM L-Glutamine (Sigma-Aldrich). For antigen-induced activation, total splenocytes isolated from OT-II mouse spleens were cultured with OVA 323-339 peptide (1 mg/ml, InvivoGen) and recombinant murine IL-2 (20 ng/ml, PeproTech). For anti-CD3/CD28-induced activation, CD4+ T cells were stimulated with Dynabeads Mouse T-Activator CD3/CD28 (Gibco) and recombinant murine IL-2 (20 ng/ml, PeproTech), following manufacturer’s instructions.

### Flow cytometry

The following antibodies were used for flow cytometry (dilution in brackets) – from BioLegend: α-CD4 BV605 GK1.5 (1:400), α-CD3 AF700 17A2 (1:200), α-CD25 PerCP-Cy5.5 PC61 (1:200), α-CD62L APC-Cy7 MEL-14 (1:400), α-CD69 PE-Cy7 H1.2F3 (1:400), α-CD127 (IL-7Rα) PE A7R34 (1:200 for surface staining, 1:100 for intracellular staining); from BD Biosciences: α-CD122 (IL-2Rβ) BV650 5H4 (1:200), α-CD132 (γc) BV650 TUGm2 (1:200); from eBioscience: α-TCRVa2 APC B20.1 (1:400), α-CD44 PE-Cy7 IM7 (1:200), α-CD8a FITC 53-6.7 (1:400); from Cell Signalling: α-phosphor-Stat5 (Tyr694) AF647 C71E5 (1:100); from Luminex: α-LC3 FITC 4E12 (1:20).

To assess the surface receptor level, cells were washed with ice-cold phosphate buffer saline (PBS) and incubated for 20 min at 4 °C in the dark, with cold PBS containing Fc block (0.125 g/ml Biolegend) and antibodies. After washing with cold PBS, cells were fixed for 10 min at room temperature with 2% Paraformaldehyde and analysed using LSRFortessa™ X-20 Cell Analyzer (BD Biosciences), LSR II Flow Cytometer (BD Biosciences) and Flowjo software (Tree Star). For intracellular staining of IL-7Rα, cells were fixed with Fixation Buffer (BioLegend) in the dark for 20 min at room temperature after surface staining, then permeabilised with Intracellular Staining Perm Wash Buffer (BioLegend) and incubated with anti-IL-7Rα antibody 1:100 diluted in Perm/Wash Buffer in the dark for 1 hour at room temperature. Cells were extensively washed with Perm/Wash buffer and resuspended with FACS buffer for further flow cytometry analysis.

To measure cell proliferation profile in vitro, cells isolated from mouse spleens were stained with CellTrace Violet (5 μM, Invitrogen) for 20 min at room temperature in dark. After extensive washing, cells were counted and seeded for activation.

To identify viable cells, cells were stained with LIVE/DEAD Fixable Aqua Stain Kit (1:400, Invitrogen) or LIVE/DEAD Fixable Near-IR Stain Kit (1:1,000, Invitrogen) diluted with PBS at 4 °C for 20 min. For apoptosis detection, splenocytes were stained with surface antibody and LIVE/DEAD Fixable Aqua Stain Kit. The cells were stained with Annexin V APC (BioLegend) 1:50 diluted in Annexin V binding buffer (BioLegend). Apoptotic cells were gated on the LIVE/DEAD-negative, Annexin V-positive population.

Tetramer staining was performed similarly as previously described^4^. In brief, before surface staining, cells were incubated with 20ug/ml tetramer in PBS at 37 °C for 30 min.

For assessing the phosphor-Stat5 level, cells were fixed with pre-heated 2% PFA for 15 min at 37 °C after the live-dead staining, permeabilised with pre-cooled 90% methanol for more than 4 hours, and stained overnight with the phospho-Stat5 antibody diluted with 0.5% BSA in PBS at 4 °C overnight.

To measure the autophagic flux, cells were treated with Bafilomycin A1 (10 nM) or DMSO for 2 hours before staining. We adapted the Guava Autophagy LC3 Antibody-based Detection Kit (Luminex) as follows: After surface staining, cells were washed once with Assay Buffer. After permeabilization with 0.05% Saponin, cells were spun immediately and incubated with anti-LC3 (FITC) antibody (1:20 diluted in Assay Buffer) at room temperature for 30 minutes in dark. After extensive washing with Assay Buffer, cells were fixed with 2% PFA and underwent flow cytometry analysis. Autophagic flux was calculated as LC3-II mean fluorescence intensity of (BafA1-Vehicle)/Vehicle.

### IL-7Rα internalisation assay

IL-7Rα internalisation assay was performed similar as described before^5^. In brief, CD4+ T cells were purified from the spleens of wild-type or T-Atg7^−/−^ mice. After incubation with biotin-conjugated anti-IL-7Rα antibody (BioLegend, A7R34) 1:50 diluted in DPBS on ice for 30 min, cells were extensively washed and fractionated into the following groups (5 × 10^6^ cells/group): 90 min on ice, 90, 60 and 30 min at 37 °C, 5% CO_2_. After incubation, cells were stained with AF647-Streptavidin (BioLegend), α-CD4 BV605 GK1.5 (BioLegend, 1:400), anti-CD44 BV785 IM7 (BioLegend, 1:400), LIVE/DEAD Fixable Aqua (Invitrogen) stain for 20 min on ice and fixed with 2% PFA. The % internalised IL-7Rα is calculated as the following equation (MFI: mean fluorescence intensity):

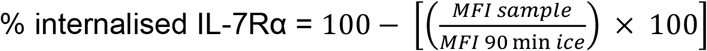

### Western blots

Cells were washed with cold PBS and lysed using RIPA lysis buffer (Sigma-Aldrich) supplemented with complete Protease Inhibitor Cocktail (Roche) and PhosSTOP (Roche). Total protein concentration in supernatant after spinning down the debris was quantified by BCA Assay (Thermo Fisher). Samples were added with 4x Laemmli Sample Buffer (Bio-Rad) and boiled at 100 °C for 5 min, with 20-30 μg protein per sample for SDS-PAGE analysis. 4–20% Mini-PROTEAN TGX Precast Protein Gels (Bio-Rad) with Tris/Glycine/SDS running buffer (Bio-Rad) was used. PVDF membrane (Millipore) transferred with protein were blocked with 5% skimmed milk-TBST. Membranes were incubated overnight with following primary antibodies diluted in 1% skimmed milk: LC3B (2775, Cell Signalling, 1:1000), APEX (IgG2A) (custom made, Regina Feederle, 1:200), GAPDH (MAB374, Millipore, 1:10,000), β-actin (3700, Cell Signalling, 1:5,000), Atg9a (ab108338, Abcam, 1:1,000), followed by 1-hour room incubation at room temperature with secondary antibodies diluted in 1% milk with 0.01% SDS: IRDye 680LT Goat anti-Mouse IgG (H + L) (Licor, 1:7,500), IRDye 680LT Goat anti-Rat IgG (H + L) (Licor, 1:10,000) and IRDye 800CW Goat anti-Rabbit IgG (H + L) (Licor, 1:10,000). Biotinylated proteins were detected by IRDye 680RD Streptavidin (Licor, 1:1,000) diluted in 1% milk with 0.01% SDS. Membranes were imaged with the Odyssey CLx Imaging System (Licor). Data were analysed using Image Studio Lite (Licor).

### Immunofluorescence staining and confocal microscopy

For intracellular staining of imMEFs, cells were seeded on coverslips in tissue culture plates. After reaching 70% confluency, proximity labelled imMEFs were washed with ice-cold PBS and fixed with 2% paraformaldehyde (Invitrogen) for 10 min, then permeabilised with 0.5% Triton × for 15 min and blocked with PBS containing 0.5% bovine serum albumin (Sigma-Aldrich) for 1 h at room temperature.

For intracellular staining of purified CD4+ T lymphocytes, cells were washed in PBS and transferred on PolyL-Lysine (Sigma-Aldrich) treated coverslips, followed by incubation for 30 min at 37 °C. For intracellular staining, cells were fixed with 2% paraformaldehyde (Sigma-Aldrich) for 10 min and permeabilised with 0.1% Triton × (Sigma-Aldrich) for 10 min, then blocked in PBS containing 2% bovine serum albumin (Sigma-Aldrich) and 0.01% Tween 20 (Sigma-Aldrich) for 1 h at room temperature.

For surface staining of purified CD4+ T lymphocytes, activated cells were washed in PBS and transferred on PolyL-Lysine (Sigma-Aldrich) treated coverslips, followed by incubation for 30 min at 37 °C. After cytokine treatment for 15 min, cells were washed and stained with antibodies at 4 °C for 20 min and fixed with 2% PFA.

All antibodies for immunofluorescence stain are as follow: FITC-LAMP1 (121605, BioLegend, 1:100), WIPI2 (SAB4200399, Sigma, manually conjugated with AF568, 1:100), AF647-Streptavidin (405237, BioLegend, 1:1,000), LC3 (PM036, MBL International, manually conjugated with AF488, 1:100), AP2 (custom made from Regina Feederle, manually conjugated with AF568), IL-7Rα (135002, BioLegend, manually conjugated with AF488, 1:100), IL-2Rα (154202, BioLegend, manually conjugated with AF568, 1:100), γc (132307, BioLegend, manually conjugated with AF647, 1:100).

After stained with antibodies for 45 min at room temperature and washed, samples were stained with DAPI (1 μg/ml, Thermo Fisher), mounted to slides with SlowFade Gold Antifade Mountant (Invitrogen), and imaged with Zeiss LSM 980 confocal microscope 63x oil immersion-lens (Zeiss). Data were analysed with ZEN Blue (Zeiss) and ImageJ software. Pearson correlation coefficients were obtained using JACoP pug-in^6^.

### Supported Lipid Bilayer (SLB) preparation and use

Glass coverslips were plasma cleaned and mounted onto six-channel chambers (Ibidi). SLB were prepared as previously described^7^, using 1,2-dioleoyl-sn-glycero-3-[(N-(5-amino-1-carboxypentyl) iminodiacetic acid) succinyl]-Ni (NTA-lipids, Avanti Polar Lipids), biotinylated 1,2-dioleoyl-sn-glycero-3-phosphoethanolamine (biotinyl cap PE-lipids, Avanti Polar Lipids) and 1,2-dioleoyl-sn-glycero-3-phosphocholine (DOPC-lipids, Avanti Polar Lipids). Channels in Ibidi chamber were covered with liposome mixture and after a 20 min incubation at room temperature, they were washed and blocked with 3% Bovine Serum Albumin (BSA, Sigma-Aldrich) and 100 μM NiSO4 for 20 min. After incubation with 10 μg/ml streptavidin, bilayers were incubated with mono-biotinylated 2c11 (anti-mouse CD3 antibody) and his-tagged mouse ICAM1-AF405, to reach a molecular density of 200 molecules/μm^2^. Channels were washed and cells were added on the bilayer and incubated at 37°C for 10 min before fixation with 4% PFA. Next, samples were permeabilized with 0.1% Saponin for 10 min and blocked with 100 mM Gly 3% BSA for 30 min. Finally, samples were incubated with primary antibodies in 1% BSA.

### Total Internal Reflection Fluorescence Microscopy (TIRFM) and image analysis

Imaging was performed on an Olympus IX83 inverted microscope equipped with a TIRF module. The instrument was equipped with an Olympus UApON 150 × 1.45 NA objective, 4-line illumination system (405, 488, 561 and 640 nm laser) and Photomertrics Evolve delta EMCCD camera. Image analysis and visualization was performed using ImageJ software. Pearson correlation coefficients were obtained using JACoP pug-in^6^.

### Electron Microscopy

*Lc3b-AP2*-expressing imMEFs were grown on aclar sheets (Science Services), supplemented with 4.83 μM Hemin chloride (ROTH) for 16 h and treated with 200 nM BafA1 (Biomol) 2 h before fixation. Cells were fixed in 2.5% glutaraldehyde (EM-grade, Science Services) in 0.1 M sodium cacodylate buffer (pH 7.4; CB) for 30 min. Fixation and the following processing steps were carried out on ice. After washes in CB, endogenous peroxidases were blocked in 20 mM glycine (Sigma) in CB for 5 min and cells washed in CB. 1x diaminobenzidine (DAB) in CB with 2 mM calcium chloride was prepared from a 10x DAB stock (Sigma) in hydrochloric acid (Sigma) and added to the cells for 5 min without and for another 40 min with 10 mM H_2_O_2_ (Sigma). After washes in CB, cells were postfixed in reduced osmium (1.15% osmium tetroxide, Science Services; 1.5% potassium ferricyanide, Sigma) for 30 min, washed in CB and water and incubated over-night in 0.5% aqueous uranylacetate (ScienceServices). Dehydration was accomplished using a graded series of ice-cold ethanol. Cell monolayers were infiltrated in epon (Serva) and cured for 48 h at 60°C. Cells were ultrathin sectioned at 50 nm on formvar-coated copper grids (Plano). TEM images were acquired on a JEM 1400plus (JEOL) using the TEMCenter and Shotmeister software packages (JEOL) and analyzed in Fiji.

### Proximity labelling

The proximity labelling based on AP2 peroxidase was performed as described before8. After 30 min biotin-phenol (500 μM, Iris Biotech) treatment at 37 °C, cells were supplemented with H_2_O_2_ (1 mM, Sigma-Aldrich) for 1 min and immediately quenched three times with cold quenching solution (10 mM sodium ascorbate, 5mM Trolox and 1mM sodium azide in DPBS). After 3 times more wash with PBS, the dry pellets of the harvested cells were flash frozen and stored in −80 °C.

### Proteinase K digestion

All steps were performed essential as previously described at 4 °C or on ice unless specified^9^. The pellets of labelled cells were washed and re-suspended in Homogenisation Buffer I (10 mM KCl, 1.5 mM MgCl_2_, 10 mM HEPES-KOH and 1 mM DTT adjusted to pH 7.5). After incubating on overhead shaker for 20 min, cell suspension was homogenised with a tight-fitting pestle in Dounce homogeniser (Scientific Laboratory Supplies) with 70 strikes and balances with 1/5 volume of Homogenisation Buffer II (375 mM KCl, 22.5 mM MgCl_2_, 220 mM HEPES-KOH and 0.5 mM DTT adjusted to pH 7.5). After centrifugation with 600x g for 10 min, clear supernatant containing autophagosomes were transferred to new tubes. For Proteinase K protection assay, supernatant was equally divided into four portions, added with 0.2% Triton X-100 (Merck) and/or 30 μg/mL proteinase K (Roche), or nothing. Following 30 min incubation at 37 °C, PMSF (10mM, Sigma-Aldrich) was administrated to inhibit the proteinase K activity. For mass spectrometry, samples were subjected to 100 μg/mL proteinase K digestion for 1 hour at 37 °C followed by PMSF treatment. For control samples 0.1% RAPIGestTM was additionally added to the proteinase K digest. Membrane-protected material was enriched by centrifugation at 17,000x g for 15 min.

### Streptavidin-pulldown and on-beads peptide digestion

The pellets of membrane-protected material were lysed with RIPA buffer containing quenchers (50 mM Tris, 150 mM NaCl, 0.1% SDS, 1% Triton X-100, 0.5% sodium deoxycholate, 1x cOmplete Protease Inhibitor Cocktail (Roche), 1x PhosSTOP (Roche), 10 mM sodium ascorbate, 1 mM Trolox and 1 mM sodium azide), sonicated and centrifuged at 10,000x g for 10 min. The supernatant was incubated with Streptavidin-agarose (Sigma-Aldrich) overnight, which was balanced with RIPA buffer containing quenchers. After 3x wash with RIPA buffer and 3x wash with 3 M Urea dissolved in 50 mM NH_4_HCO_3_, beads were incubated TCEP (5 mM, Sigma-Aldrich) at 55 °C for 30 min and shaken at 1000x rpm. Samples were alkylated with IAA (10 mM, Sigma-Aldrich) at room temperature for 20 min and shaken at 1000x rpm, further quenched by DTT (20 mM, Sigma-Aldrich) and washed 2x with 2 M Urea dissolved in 50 mM NH_4_HCO_3_. After overnight incubation with trypsin (1 μg/20 μl beads, Promega), supernatants were collected, plus 2x washes with 2 M Urea buffer. The samples were acidified with trifluoroacetic acid (1%) and underwent vacuum centrifugation to decrease the volume. After being desalted on C18 stage tips (Thermo Scientific), peptides were reconstituted with 0.5% acetic acid for mass spectrometry analysis.

### Mass spectrometry data analysis

MaxQuant (version 1.6.10.43) were used for peak detection and quantification of proteins based on RAW data. MS spectra were searched referred to the manually-annotated UniProt Mus musculus proteome (retrieved 30/03/2020), using the Andromeda search engine with the parameters as follow: full tryptic specificity, allowing two missed cleavage sites, modifications included carbamidomethyl (C) and the variable modification to acetylation (protein N terminus) and oxidation (M), and filtered with a false discovery rate (FDR) of 0.01. Analysis of label-free quantification intensities of proteins were log2-transformed with Python programming (Version 3.7.6). Missing values were replaced by random values from a distribution a quarter of the width, and −1.8 units of the original sample distributions. Proteins without greater-than-background values in both replicates for at least one condition were removed. Volcano plots were generated using GraphPad Prism software (Version 8.2.1). The log2(BafA1:DMSO) fold change of each protein is plotted on × versus the log10(p-value) of each protein plotted on y. Gene ontology analysis was performed with g:Profiler^10^.

### Statistics

Data were displayed as mean ± SEM in bar graphs, or mean ± max/min value in box plots. p-values were determined by two-tailed Student’s t-test with GraphPad Prism software (Version 8.2.1), with significant statistical differences displayed in the figure legends or graphs. For meta-analysis, proteins that were detected in both MS studies were meta-analysed using a Z-score meta-analysis weighted by the square root of sample size (N=8 for the first experiment, N=6 for the second), and the Benjamini-Hochberg procedure was used to identify significantly differential abundant proteins with FDR < 0.05.

## Ethics statement

Animal experiments were performed in accordance with institutional policies, British federal regulations, and were approved by a local ethical review committee in UK (project license PPL 30/3388).

## Acknowledgements

We thank Jonathan Webber for assistance in flow cytometry and cell sorting, Ryan Beveridge in producing lentivirus, Helena Coker for assistance in confocal microscopy, Svenja Hester for assistance in mass spectrometry, Najmeeyah Brown, Patricia Moreira and Daniel Andrew for assistance in mouse colony management. We thank Michael L. Dustin, Mark Coles, Roman Fisher, Ricardo Fernandes, Raymond Moniz and Anish Suri for scientific discussions. This work was supported by grants from the Wellcome Trust Investigator award 103830/Z/14/Z and 220784/Z/20/Z to A.K.S., Deutsche Forschungsgemeinschaft (DFG, German Research Foundation) within the frameworks of the Munich Cluster for Systems Neurology (EXC 2145 SyNergy – ID 390857198) and the Collaborative Research Center 1177 (ID 259130777) to C.B., the Wellcome Trust Investigator award 208750/Z/17/Z to L. J., the Kennedy-Chinese Scholarship Council PhD studentship to D.Z., the European Union’s Horizon 2020 (under the Marie Sklodowska-Curie grant agreement number 893676) to M.B., the Cue Biopharma Post-doctoral Fellowship, KTRR Cell Dynamics Platform and Wellcome 100262Z/12/Z to J.C.. For the purpose of Open Access, the author has applied a CC BY public copyright licence to any *Author Accepted Manuscript* version arising from this submission.

## Authors Contribution

D.Z., C.B., D.P. and A.K.S. conceptualised the study. D.Z., S.Z., C.B., L. J. and A.K.S. devised the methodology. D.Z., G.A., M.B., S.Z., J.C., D.P. and S.S. carried out the experiments. D.Z., G.A., J.C., L.J. and A.K.S. analysed and/or interpreted the experimental data. D.Z. and A.K.S. wrote the original draft. M.B., G.A., J.C., C.B. and A.K.S. reviewed and edited the manuscript. L.J., C.B., G.A. and A.K.S. provided the supervision.

## References

1 Komatsu, M. et al. Impairment of starvation-induced and constitutive autophagy in Atg7-deficient mice. J Cell Biol 169, 425–434, doi:10.1083/jcb.200412022 (2005).

2 Lee, P. P. et al. A critical role for Dnmt1 and DNA methylation in T cell development, function, and survival. Immunity 15, 763–774, doi:10.1016/s1074-7613(01)00227-8 (2001).

3 Durkin, M. E., Qian, X., Popescu, N. C. & Lowy, D. R. Isolation of Mouse Embryo Fibroblasts. Bio Protoc 3, doi:10.21769/bioprotoc.908 (2013).

4 Puleston, D. J. et al. Autophagy is a critical regulator of memory CD8(+) T cell formation. Elife 3, doi:10.7554/eLife.03706 (2014).

5 Keller, C. W. et al. The autophagy machinery restrains iNKT cell activation through CD1D1 internalization. Autophagy 13, 1025–1036, doi:10.1080/15548627.2017.1297907 (2017).

6 Bolte, S. & Cordelieres, F. P. A guided tour into subcellular colocalization analysis in light microscopy. J Microsc 224, 213–232, doi:10.1111/j.1365-2818.2006.01706.x (2006).

7 Choudhuri, K. et al. Polarized release of T-cell-receptor-enriched microvesicles at the immunological synapse. Nature 507, 118–123, doi:10.1038/nature12951 (2014).

8 Le Guerroue, F. et al. Autophagosomal Content Profiling Reveals an LC3C-Dependent Piecemeal Mitophagy Pathway. Mol Cell 68, 786–796 e786, doi:10.1016/j.molcel.2017.10.029 (2017).

9 Zellner, S., Nalbach, K. & Behrends, C. Autophagosome content profiling using proximity biotinylation proteomics coupled to protease digestion in mammalian cells. STAR Protoc 2, 100506, doi:10.1016/j.xpro.2021.100506 (2021).

10 Raudvere, U. et al. g:Profiler: a web server for functional enrichment analysis and conversions of gene lists (2019 update). Nucleic Acids Res 47, W191–W198, doi:10.1093/nar/gkz369 (2019).

11 Clarke, A. J. & Simon, A. K. Autophagy in the renewal, differentiation and homeostasis of immune cells. Nat Rev Immunol 19, 170–183, doi:10.1038/s41577-018-0095-2 (2019).

12 Sckisel, G. D. et al. Out-of-Sequence Signal 3 Paralyzes Primary CD4(+) T-Cell-Dependent Immunity. Immunity 43, 240–250, doi:10.1016/j.immuni.2015.06.023 (2015).

13 Howden, A. J. M. et al. Quantitative analysis of T cell proteomes and environmental sensors during T cell differentiation. Nat Immunol 20, 1542–1554, doi:10.1038/s41590-019-0495-x (2019).

14 Dowling, S. D. & Macian, F. Autophagy and T cell metabolism. Cancer Lett 419, 20–26, doi:10.1016/j.canlet.2018.01.033 (2018).

15 Murera, D. et al. CD4 T cell autophagy is integral to memory maintenance. Sci Rep 8, 5951, doi:10.1038/s41598-018-23993-0 (2018).

16 Jacquin, E. & Apetoh, L. Cell-Intrinsic Roles for Autophagy in Modulating CD4 T Cell Functions. Front Immunol 9, 1023, doi:10.3389/fimmu.2018.01023 (2018).

17 Lam, S. S. et al. Directed evolution of APEX2 for electron microscopy and proximity labeling. Nat Methods 12, 51–54, doi:10.1038/nmeth.3179 (2015).

18 Gingras, A. C., Abe, K. T. & Raught, B. Getting to know the neighborhood: using proximity-dependent biotinylation to characterize protein complexes and map organelles. Curr Opin Chem Biol 48, 44–54, doi:10.1016/j.cbpa.2018.10.017 (2019).

19 Xu, X. et al. Autophagy is essential for effector CD8(+) T cell survival and memory formation. Nat Immunol 15, 1152–1161, doi:10.1038/ni.3025 (2014).

20 Strutt, T. M., McKinstry, K. K., Kuang, Y., Bradley, L. M. & Swain, S. L. Memory CD4+ T-cell-mediated protection depends on secondary effectors that are distinct from and superior to primary effectors. Proc Natl Acad Sci U S A 109, E2551–2560, doi:10.1073/pnas.1205894109 (2012).

21 Barnden, M. J., Allison, J., Heath, W. R. & Carbone, F. R. Defective TCR expression in transgenic mice constructed using cDNA-based alpha- and beta-chain genes under the control of heterologous regulatory elements. Immunol Cell Biol 76, 34–40, doi:10.1046/j.1440-1711.1998.00709.x (1998).

22 Pua, H. H., Dzhagalov, I., Chuck, M., Mizushima, N. & He, Y. W. A critical role for the autophagy gene Atg5 in T cell survival and proliferation. J Exp Med 204, 25–31, doi:10.1084/jem.20061303 (2007).

23 Stephenson, L. M. et al. Identification of Atg5-dependent transcriptional changes and increases in mitochondrial mass in Atg5-deficient T lymphocytes. Autophagy 5, 625–635, doi:10.4161/auto.5.5.8133 (2009).

24 Hubbard, V. M. et al. Macroautophagy regulates energy metabolism during effector T cell activation. J Immunol 185, 7349–7357, doi:10.4049/jimmunol.1000576 (2010).

25 Jia, W. & He, Y. W. Temporal regulation of intracellular organelle homeostasis in T lymphocytes by autophagy. J Immunol 186, 5313–5322, doi:10.4049/jimmunol.1002404 (2011).

26 Parekh, V. V. et al. Impaired autophagy, defective T cell homeostasis, and a wasting syndrome in mice with a T cell-specific deletion of Vps34. J Immunol 190, 5086–5101, doi:10.4049/jimmunol.1202071 (2013).

27 Martinez, J. et al. Molecular characterization of LC3-associated phagocytosis reveals distinct roles for Rubicon, NOX2 and autophagy proteins. Nat Cell Biol 17, 893–906, doi:10.1038/ncb3192 (2015).

28 Yorimitsu, T. & Klionsky, D. J. Autophagy: molecular machinery for self-eating. Cell Death Differ 12 Suppl 2, 1542–1552, doi:10.1038/sj.cdd.4401765 (2005).

29 Zellner, S., Schifferer, M. & Behrends, C. Systematically defining selective autophagy receptor-specific cargo using autophagosome content profiling. Mol Cell 81, 1337–1354 e1338, doi:10.1016/j.molcel.2021.01.009 (2021).

30 Leidal, A. M. & Debnath, J. Emerging roles for the autophagy machinery in extracellular vesicle biogenesis and secretion. FASEB Bioadv 3, 377–386, doi:10.1096/fba.2020-00138 (2021).

31 Alves, N. L., van Leeuwen, E. M., Derks, I. A. & van Lier, R. A. Differential regulation of human IL-7 receptor alpha expression by IL-7 and TCR signaling. J Immunol 180, 5201–5210, doi:10.4049/jimmunol.180.8.5201 (2008).

32 Surh, C. D. & Sprent, J. Homeostasis of naive and memory T cells. Immunity 29, 848–862, doi:10.1016/j.immuni.2008.11.002 (2008).

33 Henriques, C. M., Rino, J., Nibbs, R. J., Graham, G. J. & Barata, J. T. IL-7 induces rapid clathrin-mediated internalization and JAK3-dependent degradation of IL-7Ralpha in T cells. Blood 115, 3269–3277, doi:10.1182/blood-2009-10-246876 (2010).

34 Faller, E. M., Ghazawi, F. M., Cavar, M. & MacPherson, P. A. IL-7 induces clathrin-mediated endocytosis of CD127 and subsequent degradation by the proteasome in primary human CD8 T cells. Immunol Cell Biol 94, 196–207, doi:10.1038/icb.2015.80 (2016).

35 Loi, M. et al. Macroautophagy Proteins Control MHC Class I Levels on Dendritic Cells and Shape Anti-viral CD8(+) T Cell Responses. Cell Rep 15, 1076–1087, doi:10.1016/j.celrep.2016.04.002 (2016).

36 Leonard, W. J., Lin, J. X. & O’Shea, J. J. The gammac Family of Cytokines: Basic Biology to Therapeutic Ramifications. Immunity 50, 832–850, doi:10.1016/j.immuni.2019.03.028 (2019).

37 Ross, S. H. & Cantrell, D. A. Signaling and Function of Interleukin-2 in T Lymphocytes. Annu Rev Immunol 36, 411–433, doi:10.1146/annurev-immunol-042617-053352 (2018).

38 Waickman, A. T. et al. The Cytokine Receptor IL-7Ralpha Impairs IL-2 Receptor Signaling and Constrains the In Vitro Differentiation of Foxp3(+) Treg Cells. iScience 23, 101421, doi:10.1016/j.isci.2020.101421 (2020).

39 Li, C. & Park, J. H. Assessing IL-2-Induced STAT5 Phosphorylation in Fixed, Permeabilized Foxp3(+) Treg Cells by Multiparameter Flow Cytometry. STAR Protoc 1, 100195, doi:10.1016/j.xpro.2020.100195 (2020).

40 Sabatos, C. A. et al. A synaptic basis for paracrine interleukin-2 signaling during homotypic T cell interaction. Immunity 29, 238–248, doi:10.1016/j.immuni.2008.05.017 (2008).

41 Arata, Y. et al. Defective induction of the proteasome associated with T-cell receptor signaling underlies T-cell senescence. Genes Cells 24, 801–813, doi:10.1111/gtc.12728 (2019).

42 McLeod, I. X., Zhou, X., Li, Q. J., Wang, F. & He, Y. W. The class III kinase Vps34 promotes T lymphocyte survival through regulating IL-7Ralpha surface expression. J Immunol 187, 5051–5061, doi:10.4049/jimmunol.1100710 (2011).

43 Willinger, T. & Flavell, R. A. Canonical autophagy dependent on the class III phosphoinositide-3 kinase Vps34 is required for naive T-cell homeostasis. Proc Natl Acad Sci U S A 109, 8670–8675, doi:10.1073/pnas.1205305109 (2012).

44 Lindqvist, L. M., Simon, A. K. & Baehrecke, E. H. Current questions and possible controversies in autophagy. Cell Death Discov 1, doi:10.1038/cddiscovery.2015.36 (2015).

45 Leidal, A. M. et al. The LC3-conjugation machinery specifies the loading of RNA-binding proteins into extracellular vesicles. Nat Cell Biol 22, 187–199, doi:10.1038/s41556-019-0450-y (2020).

46 Huber, L. A. et al. Rab8, a small GTPase involved in vesicular traffic between the TGN and the basolateral plasma membrane. J Cell Biol 123, 35–45, doi:10.1083/jcb.123.1.35 (1993).

47 Dupont, N. et al. Autophagy-based unconventional secretory pathway for extracellular delivery of IL-1beta. EMBO J 30, 4701–4711, doi:10.1038/emboj.2011.398 (2011).

48 Szatmari, Z. & Sass, M. The autophagic roles of Rab small GTPases and their upstream regulators: a review. Autophagy 10, 1154–1166, doi:10.4161/auto.29395 (2014).

49 Jia, W. et al. Autophagy regulates T lymphocyte proliferation through selective degradation of the cell-cycle inhibitor CDKN1B/p27Kip1. Autophagy 11, 2335–2345, doi:10.1080/15548627.2015.1110666 (2015).

50 Mocholi, E. et al. Autophagy Is a Tolerance-Avoidance Mechanism that Modulates TCR-Mediated Signaling and Cell Metabolism to Prevent Induction of T Cell Anergy. Cell Rep 24, 1136–1150, doi:10.1016/j.celrep.2018.06.065 (2018).

51 Dingar, D. et al. BioID identifies novel c-MYC interacting partners in cultured cells and xenograft tumors. J Proteomics 118, 95–111, doi:10.1016/j.jprot.2014.09.029 (2015).

52 Uezu, A. et al. Identification of an elaborate complex mediating postsynaptic inhibition. Science 353, 1123–1129, doi:10.1126/science.aag0821 (2016).

53 Liu, G. et al. Mechanism of adrenergic CaV1.2 stimulation revealed by proximity proteomics. Nature 577, 695–700, doi:10.1038/s41586-020-1947-z (2020).

54 Rudolph, F. et al. Deconstructing sarcomeric structure-function relations in titin-BioID knock-in mice. Nat Commun 11, 3133, doi:10.1038/s41467-020-16929-8 (2020).

55 Kim, K. E. et al. Dynamic tracking and identification of tissue-specific secretory proteins in the circulation of live mice. Nat Commun 12, 5204, doi:10.1038/s41467-021-25546-y (2021).

56 Feng, W. et al. Identifying the Cardiac Dyad Proteome In Vivo by a BioID2 Knock-In Strategy. Circulation 141, 940–942, doi:10.1161/CIRCULATIONAHA.119.043434 (2020).

