## Supplementary Figures for "Mapping of the autophagosomal degradome identifies IL-7Rα as key cargo in proliferating CD4+ T-cells"

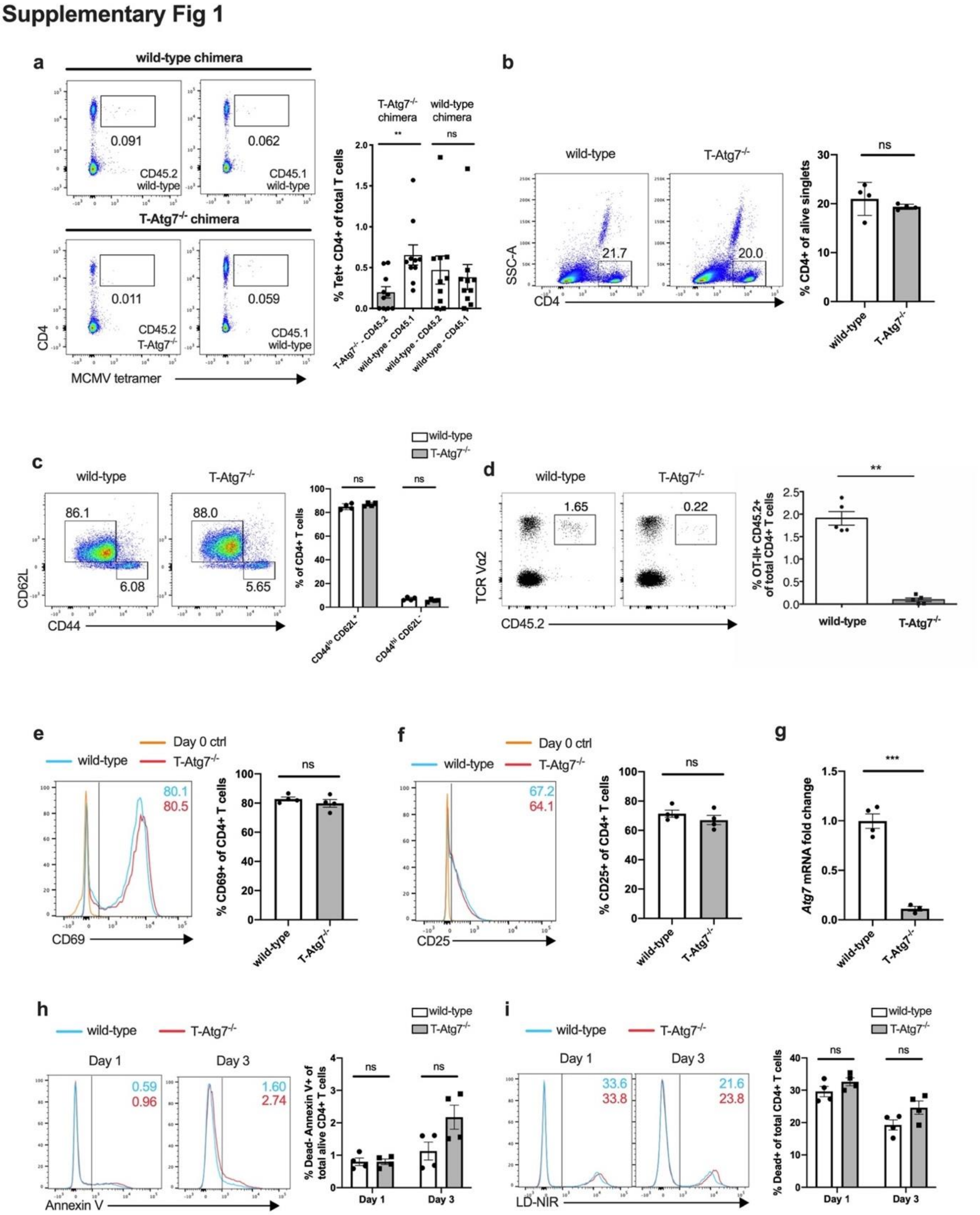

**Supplementary Figure 1. Autophagy-deficient CD4+ T cells show delayed proliferation.**

**(a)** Dot plots (left) of MCMV-specific CD4+ T cells among CD19- population from lungs of CD45.2+ wild-type or T-Atg7$^{-/-}$ donors and CD45.1+ donors. Bar graph (right) indicates the percentage of Tet+ CD4+ T cells within CD45.2+ CD19- or CD45.1+ CD19- population (n = 9 mice per group) as mean ± SEM with paired two-tailed Student’s t-test, p**<0.01, ns: not significant.

**(b, left)** and **(c, left)** show dot plots of splenic CD4+ T cells, and CD44$^{hi}$ CD62L$^{-}$ T cells within total CD4+ T cells, in 6–8-week wild-type or T-Atg7$^{-/-}$ OT-II mice. Bar graphs **(b and c, right)** indicate the percentages of gated populations within total splenocytes (n = 4 mice per group).

**(d)** Dot plots (left) are gated on CD45.2+, TCR Vα2+ in the blood of recipient mice. Bar graph (right) depicts the frequency of gated population within total CD4$^{+}$ T cell population (n = 5 mice per group).

**(e, left)** and **(f, left)** show histograms of CD69+ and CD25+ population within CD4+ T cells on day 1 of activation in vitro. Bar graphs **(e and f, right)** indicate the percentages of gated populations within live CD4+ population (n = 4 mice per group).

**(g)** Quantification of *Atg7* mRNA level in FACS-sorted most divided splenic CD4+ T cells activated for 7 days (gating strategy in Figure 1e, day 7) by qRT-PCR. Bar graph shows fold change in *Atg7* mRNA of T-Atg7$^{-/-}$ normalised to wild-type (n = 3-4 mice per group) as mean ± SEM with unpaired two-tailed Student’s t-test,.

**(h, left)** and **(i, left)** show histograms of Annexin V+ population within LD-NIR- CD4+ T cells, and LD-NIR+ population within CD4+ T cells, on day 1 and 3 of activation in vitro. Bar graph **(h and i, right)** indicates the percentage of gated population within live CD4+ population (n = 4 mice per group).

All data except for (a) are represented as mean ± SEM with unpaired two-tailed Student's t-test, $p^{**}<0.01$, $p^{***}<0.001$, ns: not significant. All experiments are representative of three independent experiments.

### Supplementary Figure 2

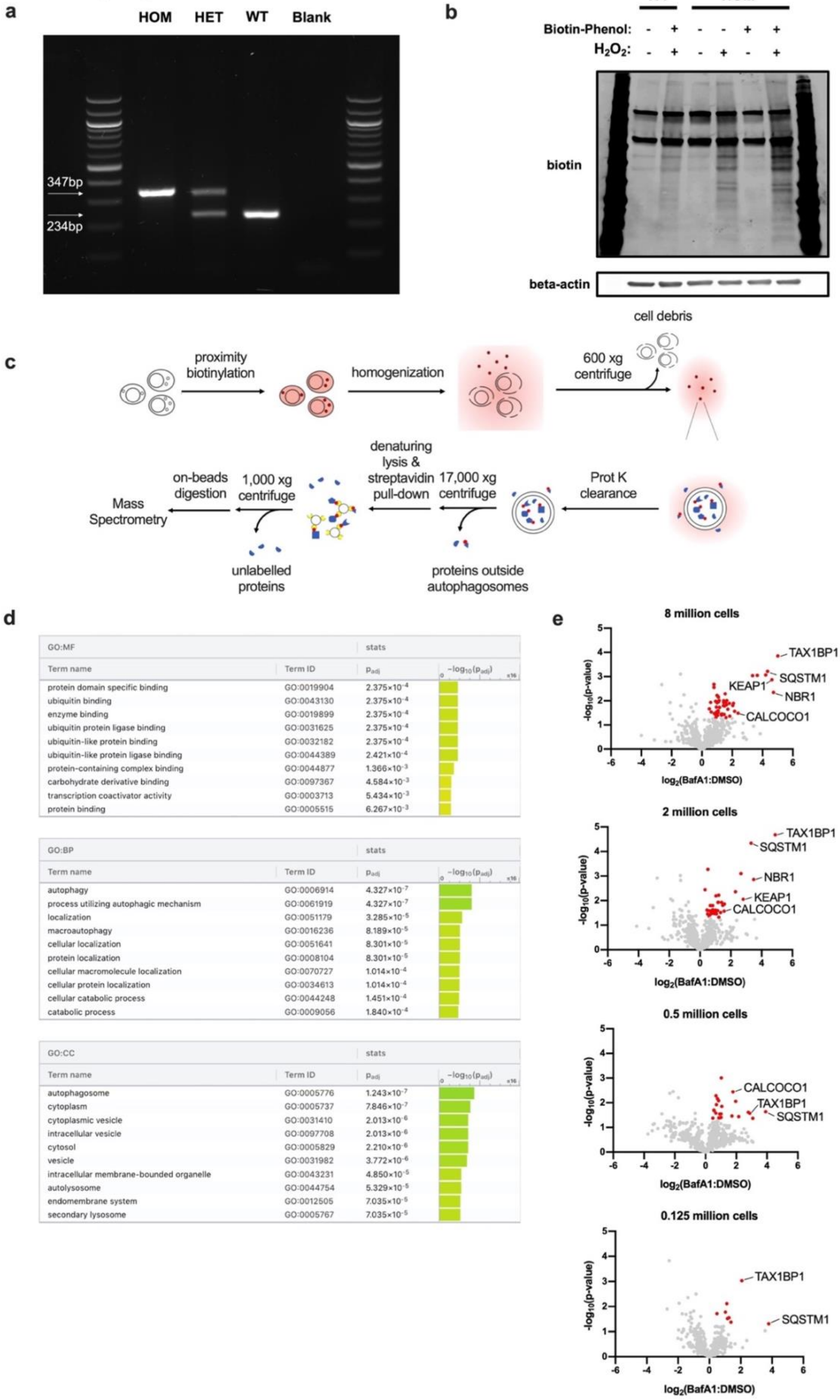

**Supplementary Figure 2. Validation of the *Lc3b-AP2* transgenic mouse model.**

**(a)** PCR genotyping of the *Lc3b-AP2* mouse model (WT, wild-type; HET, heterozygote; HOM, homozygote).

**(b)** Biotinylated protein profiles of whole-cell homogenates from splenocytes of wild-type (WT) or homozygotes (HOM), activated with 10 µg/mL LPS for 3 days. Cells were left untreated or labelled with biotin-phenol and/or $H_2O_2$. These experiments were repeated as biological triplicates with similar results.

**(c)** Schematic overview of the procedure of purifying autophagosomal biotinylated proteins.

**(d)** Gene Ontology (GO) enrichment analysis of BafA1-upregulated proteins with the top terms for molecular function (MF), biological pathway (BP) and cellular compartment (CC). $p_{adj}$ are p-values adjusted for multiple testing using the Benjamini-Hochberg method.

**(e)** Volcano plots of proteins labelled by APEX2-LC3B in immortalised MEF cells, with the titration of different cell numbers. Proteins significantly upregulated in BafA1-treated samples are highlighted in red ($p<0.05$ by unpaired two-tail Student's t-test, n = 3 biological replicates per group).

### Supplementary Figure 3

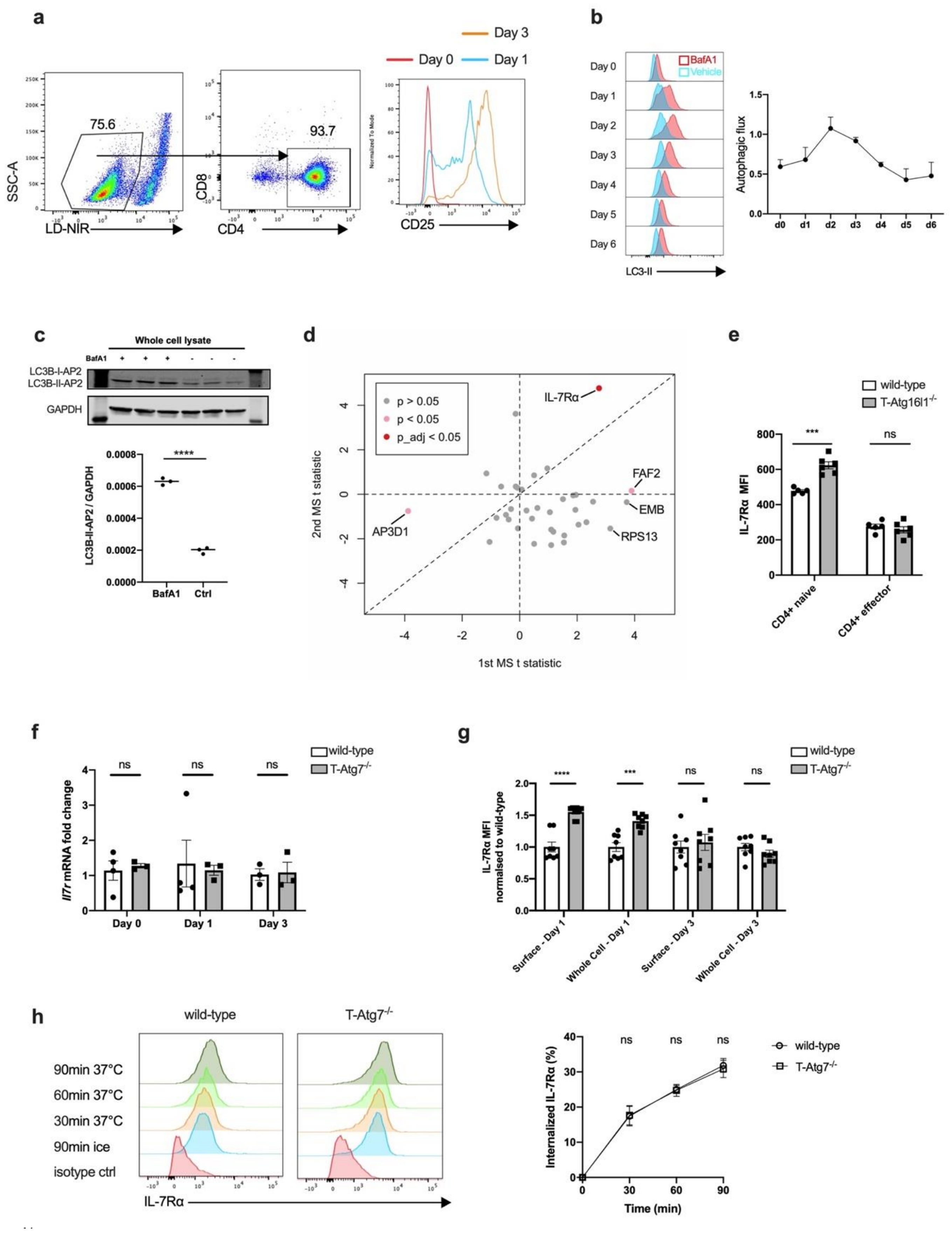

**Supplementary Figure 3. IL-7Rα is an autophagosomal cargo in CD4+ T cells.**

**(a)** Dot plots (left) displaying the survival rate and the purity of CD4+ T cells 3 days post-activation. Histogram (right) depicts the surface level of CD25 gated on live CD4+ T cells on days 0, 1 and 3 of activation.

**(b)** LC3-II histograms (left) of activated CD4+ T cells on corresponding days, which were treated or not with BafA1 for 2 hours before staining. Autophagic flux (right) was calculated by the LC3-II geometric mean fluorescent intensity (BafA1 - Vehicle)/Vehicle, n = 3 mice. Data are represented as mean ± SEM.

**(c)** Western blot (upper) and quantitative analysis (lower) of LC3B-AP2 expression level in activated CD4+ T cells on day 3 post-activation, n = 3 mice per group.

**(d)** Meta-analysis of 36 proteins detected in both proximity-biotinylation experiments in 3-day-activated CD4+ T cells. Pink dots indicate the proteins upregulated upon BafA1-treatment with nominal $p < 0.05$, whereas red dot indicates the protein with $p < 0.05$ after Benjamini-Hochberg correction.

**(e)** Bar graph depicting surface IL-7Rα level in different CD4+ subtypes from wild-type and CD4$^{cre}$ *Atg16l1*$^{fl/fl}$ (T-Atg16l1$^{-/-}$) mice, n = 5-6 mice per group.

**(f)** Quantification of *Il7ra* mRNA expression in splenic CD4+ T cells without activation, activated for 1 day or 3 days. Bar graph shows the fold change in *Il7ra* mRNA of T-Atg7$^{-/-}$ normalised to wild-type, n = 3-4 mice per group.

**(g)** Bar graph summarising flow results of both surface and whole-cell IL-7Rα level in CD4+ T cells activated for 1 or 3 days in vitro. Fold change in IL-7Rα geometric mean fluorescence intensity of T-Atg7$^{-/-}$ normalised to wild-type is shown, n = 8 mice per group.

**(h)** Internalisation of surface IL-7Rα on naïve CD4+ T cells was performed with a biotin-based flow cytometric endocytosis assay. Representative histograms (left) to show biotin-conjugated IL-7Rα antibody level. Connected scatter plot (right) indicates the percentage of internalised IL-7Rα at 30, 60, and 90 min at 37 °C, n = 6 mice per group. All values represented as mean ± SEM with unpaired two-tailed Student’s t-test, p***<0.001, p****<0.0001, ns: not significant.

### Supplementary Figure 4

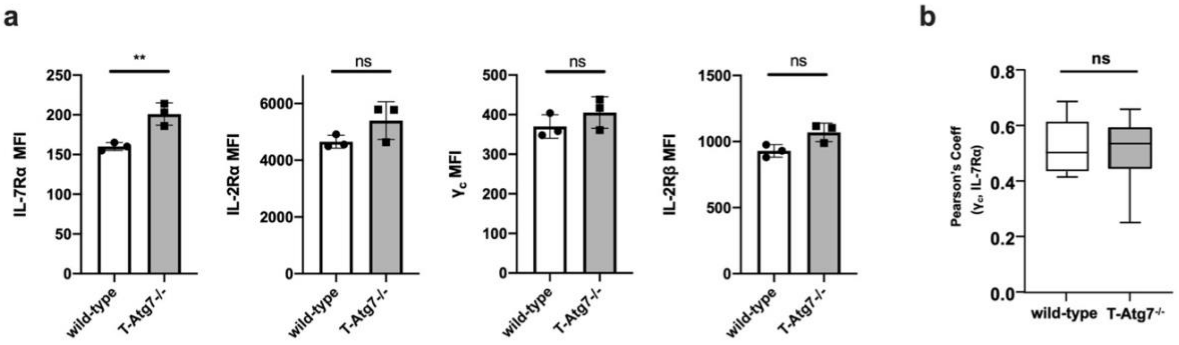

**Supplementary Figure 4. Impaired IL-2R assembly in activated T-Atg7$^{-/-}$ CD4+ T cells.**

**(a)** Splenic naïve CD4+ T cells purified from either wild-type or T-Atg7$^{-/-}$ mice were activated with anti-CD3/CD28 Dynabeads for 24 hours. The surface receptor levels were assessed with flow cytometry. Bar graphs depict surface levels of IL-7Rα, IL-2Rα, common gamma chain ($γ_c$) and IL-2Rβ on activated CD25+ CD4+ T cells, n = 3 mice per group. All data are represented as mean ± SEM with unpaired two-tailed Student's t-test, p**<0.01, ns: not significant. Quantitative analyses are representative of three independent experiments with similar results.

**(b)** Naive CD4+ T cells were pre-treated with anti-CD3/CD28 Dynabeads for 24 hours to upregulate IL-2Rα. Then, cells were exposed to supported lipid bilayers containing ICAM1 and anti-CD3 proteins and fixed after a 10-min incubation in the presence of 50 U/ml of mIL-2. Synapse formation was confirmed by the presence of an ICAM1 ring. Cytokine receptors were stained for TIRF imaging. Box graph shows Pearson's correlation coefficient of the co-localisation of IL-7Rα and $γ_c$ at the mature immune synapses of activated CD25+ CD4+ T cells, which were exposed to lipid bilayers and incubated with murine IL-2, n = 15 cells per group. Data are represented as mean ± max and min values, with unpaired two-tailed Student's t-test, ns: not significant.
